# High-Throughput Multiomics Profiling of Model Systems Using the AVITI24 Platform

**DOI:** 10.1101/2025.05.03.651997

**Authors:** Tyler Lopez, Daniel Honigfort, Adeline Mah, Connor Thompson, Vivian Dien, Mariam Dawood, Eva Frankel, Derek Fuller, Claudia Jamison, Ryan Kelley, Jean Kwon, Yu Liu, Pin Ren, Sanchari Saha, Haosen Wang, Jennifer Wong, Da Zhao, Minna Abtahi, Andrew Altomare, Rosi Bajari, Ava Bellizzi, Sierra Bracamonte, Alex Bradfield, Colin Brown-Greaves, Chris Bui, Katia Charov, Rodger Constandse, Kim Cullion, West Damron, Mike Dangelo, Laura Davis, Neaam Dawood, Rushikesh Dhonde, Keerthana Elango, Sara Espinosa, Francisco Garcia, Vlad Gavrila, Laura Gomez, Daniel Hastings, Dennis Hoang, William Juan, Amirali Kia, Michael Kim, Marielle Krivit, Roberto Lama, Kyle Mandla, Anyssa Martinez, Max Mass, Aaron Miller, Jordan Neysmith, Jesus Nunez, Pablo Radilla, Tristin Rammel, Michael Ray, Chloe Sermet, Luqmanal Sirajudeen, Ben Song, Hermes Taylor, Will Thornbury, Ramreddy Tippana, Henry Vuong, Jessica Winger, Jerry Wu, Jane Yao, Shuyang Ye, Grace Yeo, Joseph Xiao, Semyon Kruglyak, Matthew Kellinger, Michael Previte, Molly He, Sinan Arslan

## Abstract

We present a multiomics platform comprising Teton, a detection assay system, and AVITI24, a dual-flowcell instrument that performs both cellular imaging and sequencing readout. Teton integrates a compartmentalized flowcell for cell culture with methods to measure morphology, RNA, and protein at subcellular resolution. The platform quantifies morphological features through cell painting of 6 cellular components, RNA expression of up to 350 transcripts via sequencing of oligonucleotides hybridized to mRNA, and protein expression of up to 200 targets using antibody-linked oligonucleotide sequencing. The flow cell accommodates >1 million cells in a 10 cm squared open-well format or can be subdivided into 12 or 48 wells to support experiments with multiple conditions or time points. We describe and validate the detection methods of the platform and showcase its capabilities by co-culturing three cancer cell lines and elucidating the cellular pathways triggered by various drug treatments as a function of time. Using multiple time points enables us to capture the dynamics of cellular processes including receptor activation and signaling cascades. The results demonstrate how different cancer cells evade TNFα-induced apoptosis by activating compensatory signaling programs that maintain survival despite pro-apoptotic cues. Our model system replicates previously published results and highlights the versatility of the platform in enabling rapid, high-throughput analysis of complex cellular responses in varied biological contexts.

## Introduction

Cells integrate information across multiple molecular layers, from gene expression and protein signaling to morphological context, yet profiling these layers together remains a major technical challenge. Understanding heterogeneous cell states in development and disease requires analyses that capture morphology, RNA, and protein markers at single-cell resolution and at discrete time points. Traditional assays fall short: western blotting and ELISA average over populations, while microscopy-based immunofluorescence (IF) and mass spectrometry (MS) provide only partial views. IF is limited to ∼1–10 markers per assay due to spectral or host constraints, and MS requires cell lysis, losing spatial context [1,2]. Single-cell RNA sequencing (scRNA-seq) has transformed transcriptomic profiling [3,4] but measures only mRNA and requires cell dissociation. Collectively, current methods lack the ability to deliver a simultaneous, high-throughput, single-cell view of molecular content and spatial architecture [5,6]. This gap impedes our understanding of complex biological processes that are orchestrated across multiple molecular modalities in each cell.

Cell signaling networks are regulated at multiple levels, including transient protein phosphorylation and transcriptional feedback, which unfold rapidly across space and time. RNA measurements alone cannot resolve many signaling events, as protein-level changes often occur without corresponding mRNA shifts [7,8]. Many signaling events are spatially organized, beginning at membranes, junctions, or organelles and transmitting signals through the cytoplasm to the nucleus [9]. Therefore, understanding the propagation of signaling behavior in time requires multiple integrated measurements of morphology, protein states, and gene expression within the same cells [10,11,12].

Emerging technologies now support multi-modal profiling. Multiplexed cytometry and sequencing-based methods allow simultaneous RNA and protein measurement at single-cell resolution [9,13], helping to capture dynamic signaling networks [10]. However, trade-offs remain between resolution, throughput, and data integration. Droplet-based scRNA-seq (e.g., 10x Genomics Chromium) offers transcriptome-wide throughput [3] but requires cell dissociation and lacks protein or morphological data. Spatial transcriptomics methods like 10x Visium and Slide-seq preserve spatial context [14] but typically offer lower resolution or narrower coverage. In situ hybridization platforms such as NanoString’s CosMx and 10x Xenium push resolution further [9,15] but rely on iterative imaging, limited protein content, and multi-day workflows that limit scalability.

High-plex proteomic platforms also face trade-offs. Mass cytometry (CyTOF) can profile >40 proteins per cell but lacks spatial resolution and is destructive. Imaging methods like CODEX and Lunaphore COMET can reach approximately 40–60 targets per section [13], using barcoded antibodies with iterative fluorescence cycles [16,17]. These methods are affected by variability in antibody clone selection and conjugation chemistry. Moreover, few imaging platforms reliably detect phosphorylated proteins at scale. Soluble protein platforms such as Olink and SomaScan prioritize proteomic breadth: Olink detects up to 5,400 proteins per panel [18,19], and SomaScan can profile around 7,000 proteins in bulk samples [20]. However, both are limited to lysates or body fluids and lack spatial resolution, transcriptomic integration, and single-cell specificity [20].

In summary, existing technologies each excel in one area, be it throughput, spatial fidelity, or protein/RNA depth, but none fully unify all three or enable coordinated temporal measurements across modalities. A platform capable of simultaneously capturing the dynamic temporal profiles of single-cell morphology, RNA, and protein (including phosphoproteins) at scale remains an unmet need in the field.

Apoptosis provides a clear example of the complexity that demands integrated, dynamic measurement. The TNFα signaling axis can activate both apoptotic and survival pathways. TNFα binding to TNFR1 triggers extrinsic apoptosis via caspase-8, but many cells resist through NF-κB, MAPK, and Akt signaling to avoid death [21,22]. For instance, phosphorylation of heat shock protein 27 (HSP27) enhances cell survival by stabilizing the cytoskeleton and inhibiting caspase activation, cooperating with anti-apoptotic BCL-2 family proteins to block apoptosis [23,24,25,26]. The MAPK family kinases further illustrate pathway cross-talk: p38 MAPK normally promotes apoptosis in response to stress, but its inhibition can trigger compensatory activation of ERK1/2 signaling, thereby driving cell proliferation and drug resistance [23,27,28]. Even JNK, typically pro-apoptotic, may either facilitate or impede cell death depending on its interplay with NF-κB and Akt pathways [21,27, 29,30]. In cancer, such redundant and feedback-laden circuitry enables therapy resistance. Many tumors evade apoptosis by disrupting regulatory checkpoints. For example, TP53 mutations disable p53-mediated cell death, while PTEN loss activates the PI3K-Akt survival axis, promoting resistance to targeted therapies [22,31]. The TRAIL receptor pathway is similarly modulated in cancer, where pro-apoptotic signals are frequently bypassed through adaptive rewiring [32,33,34,35]. Understanding these escape mechanisms is critical for therapeutic development and underscores the need for platforms that can holistically track protein activation, gene expression, and phenotypic outcomes at single-cell resolution.

To overcome limitations in current multiomic technologies, we developed Teton on the AVITI24 platform—a high-throughput, single-cell system that simultaneously measures RNA, protein, and morphology at subcellular resolution. This integrated approach eliminates tradeoffs between spatial resolution and throughput while bridging the gap between morphological and molecular analyses. Cells cultured directly on a custom flowcell undergo an automated 24-hour workflow that extracts morphological features through high-content imaging (similar to Cell Painting [36]) while quantifying up to 350 RNA targets using barcoded hybridization probes and 200 protein targets, including phosphoproteins, via DNA-barcoded antibodies. With its ∼20 cm^2^ imaging area accommodating millions of cells per dual-sided run, Teton achieves droplet-based throughput while preserving spatial context. This creates comprehensive datasets linking each cell’s morphology to its transcriptomic and proteomic profiles at specific timepoints. We first validate Teton detection through comparison to independent technologies and then demonstrate its capabilities using a TNFα stimulation model with kinase inhibitors, revealing how this platform enables insights into molecular signaling networks, drug responses, phenotypic heterogeneity, and temporal dynamics within a unified framework.

## Results

### Teton on AVITI24 Platform Overview

Figure 1 illustrates the detection components of the Teton assay and workflow on AVITI24 which proceeds as follows. Cells were first cultured and seeded into flowcells, then fixed, labeled with cell paint probes, and permeabilized to allow for downstream molecular detection. The assembled flowcell was loaded onto the AVITI24 system for simultaneous detection of DNA-barcoded antibodies and RNA probes. Protein probes were first bound to their targets, followed by RNA probe hybridization to their corresponding transcripts. These barcoded probes were then co-amplified via rolling circle amplification to form polonies, which are highly localized concatemers of amplified DNA within individual cells.

**Figure 1:**
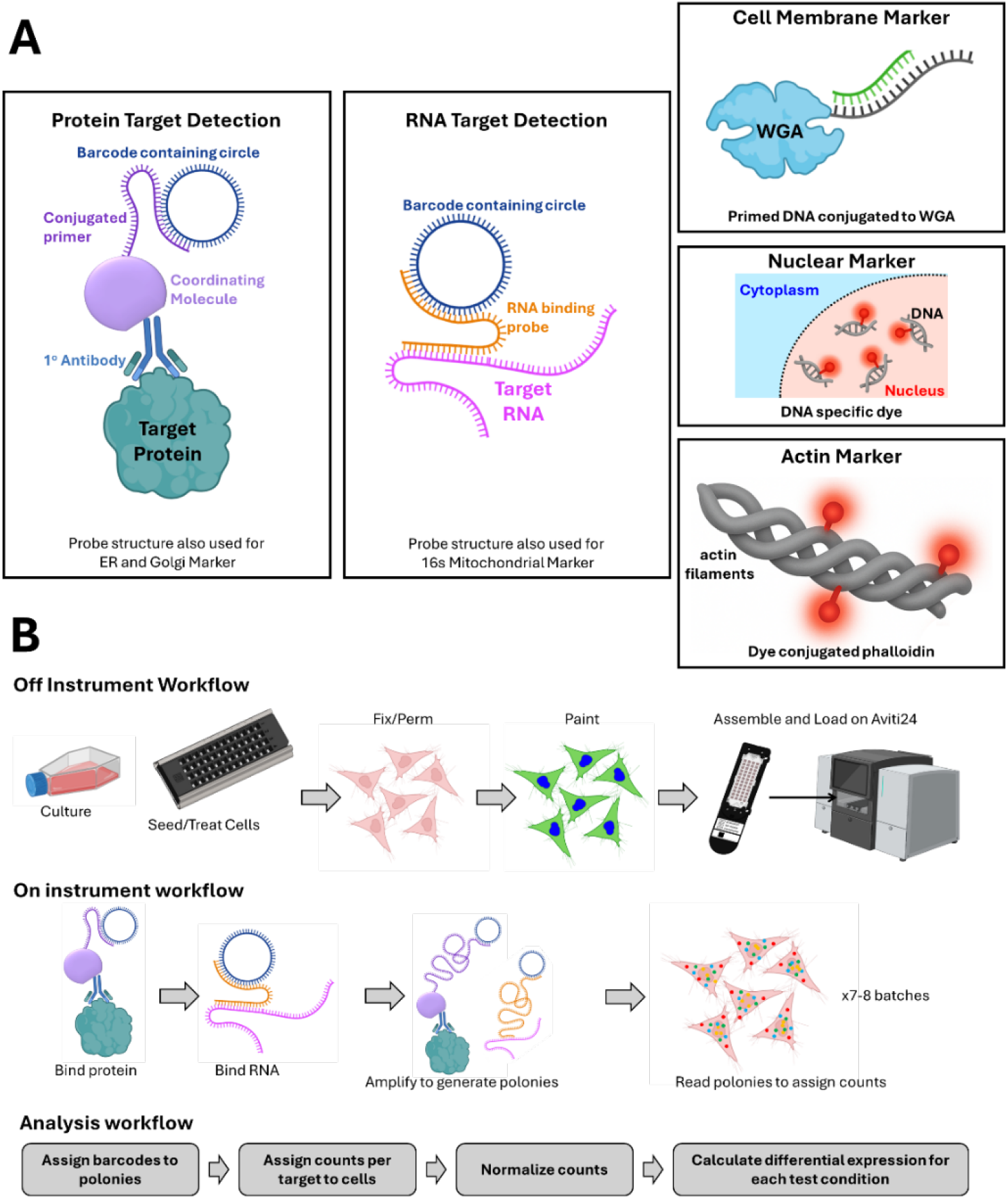
Integrated Teton detection chemistry and workflow of the AVITI24 platform. Top panels illustrate the three detection strategies used by Teton. Protein targets are detected using a primary antibody complexed with a noncovalent coordinating molecule linked to a primer and circular DNA containing a barcode; this structure is also used for detecting cell painting targets such as ER and Golgi. RNA targets are hybridized with sequence-specific probes and barcoded circular DNA for amplification and quantification; this probe design is also used to detect 16S mitochondrial RNA. Cell segmentation is performed using morphological markers specific to membrane (WGA), nucleus (RedDot2), and actin (phalloidin), which are labeled with fluorescent or DNA-conjugated reagents. Bottom panels depict the high-throughput Teton workflow on AVITI24. Cells are seeded directly onto the flowcell, stimulated, fixed, and incubated with a combination of barcoded and non-barcoded probes. Rolling circle amplification generates DNA polonies from barcoded targets, which are sequenced and assigned to single cells, enabling target quantification, normalization, and differential expression analysis. Parts of images were created using Biorender.com.

Each polony contained a unique barcode corresponding to a specific RNA or protein target. Following imaging, barcodes were decoded using standard avidite-based sequencing [37] and assigned to individual cells through spatial registration. The resulting dataset provided counts for each target on a per-cell basis, which were then normalized and used to calculate differential expression relative to the untreated control condition. This enabled robust differential expression analysis across both RNA and protein modalities. Teton used a subsampling-based detection strategy by controlling the concentration of probes rather than saturating all available target molecules. By design, only a fraction of molecules per target were amplified and sequenced, avoiding signal crowding and preserving high spatial resolution. We confirmed that this subsampling approach yielded highly reproducible target quantification across replicate wells and treatment conditions, ensuring quantitative accuracy without compromising sensitivity (Supplemental Figure 2). In addition to subsampling, Teton employed a batching strategy where only a subset of targets was read per batch. This was achieved by hybridizing a sequencing primer specific to the targets within the designated batch, reading the barcode via sequencing, dehybridizing, and repeating the process for the subsequent batch. This provided a second layer of protection against signal saturation within a single cell.

### Protein Quantification

To validate the fidelity of the Teton assay, we performed independent assays to confirm the accuracy of both protein and RNA detection. Antibody-based probes designed to bind unique circular DNA barcodes (Figure 1) were validated through immunofluorescence and western blotting across multiple cell types. Phospho-HSP27 regulation was confirmed using Teton, immunofluorescence, and western blotting, with all three methods detecting strong TNFα-induced upregulation and complete suppression upon p38 inhibition with SB202190 (SB) (Figure 2).

**Figure 2:**
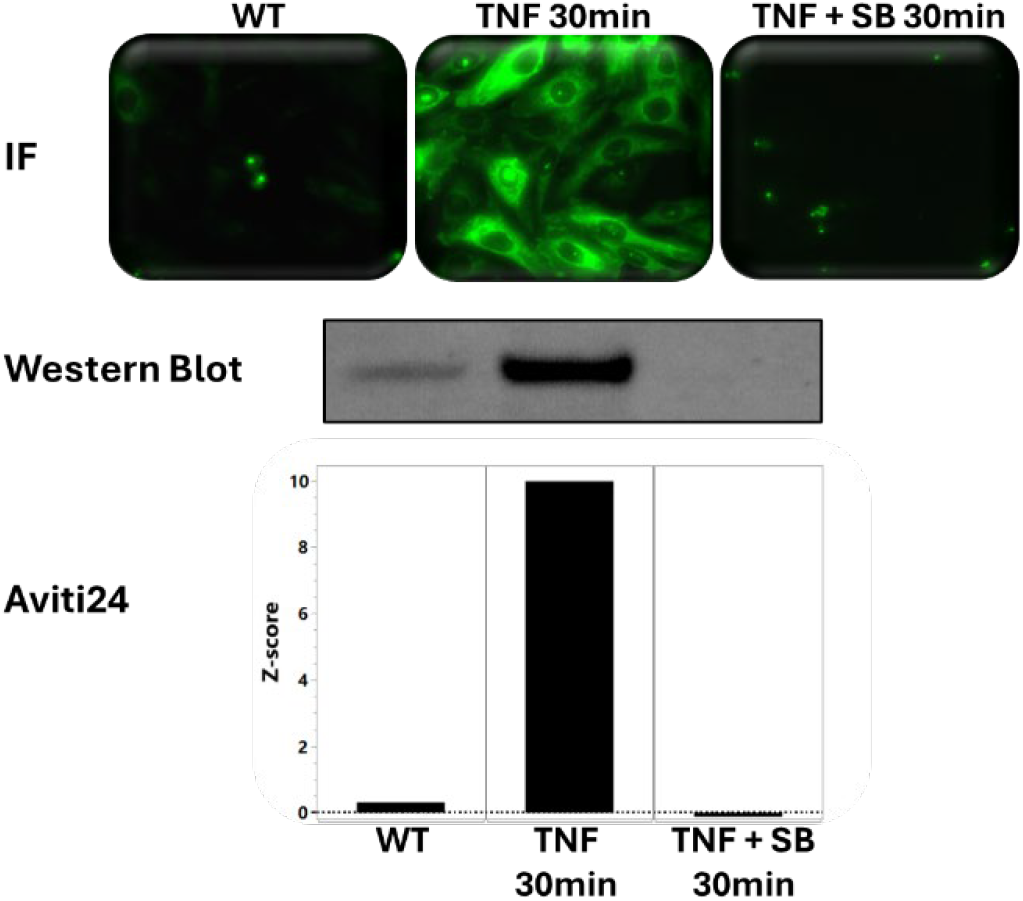
Concordant regulation of phospho-HSP27 across Teton on AVITI24, western blotting, and immunofluorescence. Phospho-HSP27 levels were measured in HeLa cells following TNFα stimulation (30 min) and p38 inhibition (TNF + SB). All three methods (AVITI24 barplot, western blot gel, and immunofluorescence images) showed strong upregulation of phospho-HSP27 with TNFα treatment and complete suppression upon addition of the p38 inhibitor SB202190 (SB).

Additional protein targets profiled across HeLa, A549, and HepG2 cells showed consistent expression trends between Teton and western blot analysis (Supplemental Figure 3), further supporting the platform’s quantitative accuracy across diverse proteins. Subsequent validation came from phosphatase treatments, which confirmed probe specificity for multiple phosphoproteins, and siRNA knockdowns of HSP60, ERK1, and ERK2, further reinforcing target specificity (Figure 3, Supplemental Figure 4).

**Figure 3:**
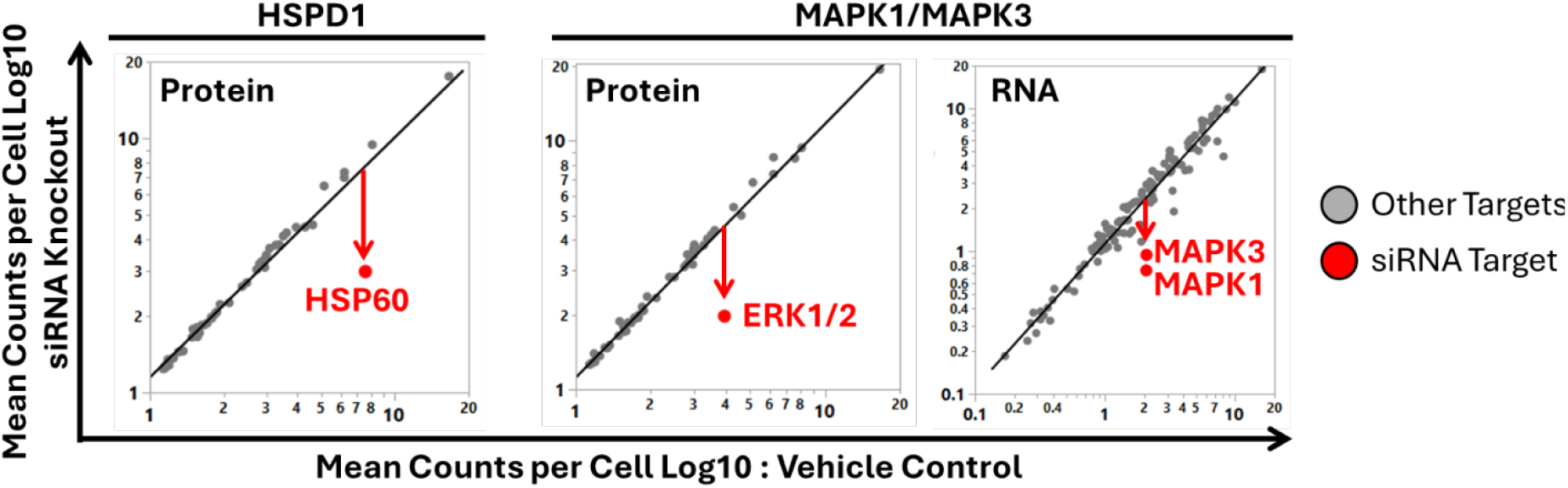
Teton detects reduced target expression following siRNA knockdown. HeLa cells were transfected with siRNAs targeting HSPD1 (HSP60), MAPK3 (ERK1), or MAPK1 (ERK2), and protein levels were measured using the Teton assay. Significant decreases in Teton signal were observed for each targeted protein, confirming the sensitivity and specificity of the antibody-barcode detection strategy. Corresponding immunofluorescence images showing knockdown efficiency are provided in Supplemental Figure 4.

To benchmark Teton against an orthogonal proteomics method, we submitted matched samples for label-free mass spectrometry analysis. Mass spec identified approximately 5,800 proteins and 52,000 peptides across all conditions, consistent with expected global coverage. However, differential expression analysis between technical replicates (Hela_WT1 and Hela_WT2) identified 1,220 proteins. Despite a high correlation (R^2^ = 0.905) between the replicates, individual targets can have orders of magnitude differences in technical replicates making it difficult to extract a meaningful conclusion from these comparisons. (Supplemental Figure 5). In contrast, Teton measurements showed no differential expression between technical replicates.

### mRNA Quantification

Short complementary DNA strands annealed with unique circular DNA barcodes were designed to detect RNA targets in cells (Figure 1). RNA detection was validated through RNA-seq, single-cell RNA-seq (scRNA-seq), and fluorescence in situ hybridization (FISH), with correlation coefficients of R^2^ > 0.67 for bulk RNA-seq, R^2^ > 0.72 for scRNA-seq, and R^2^ > 0.93 for FISH (Figure 4). The higher correlation observed with FISH may reflect its direct digital readout, in contrast to the amplification-based workflows used in RNA-seq and scRNA-seq. Ongoing studies aim to further investigate this observation.

By integrating multiple validation methods, including RNA-seq, scRNA-seq, FISH, immunofluorescence, mass spectrometry, western blotting, and siRNA knockdowns, we confirmed that Teton measurements are consistent with independent methods for both protein and RNA targets.

**Figure 4:**
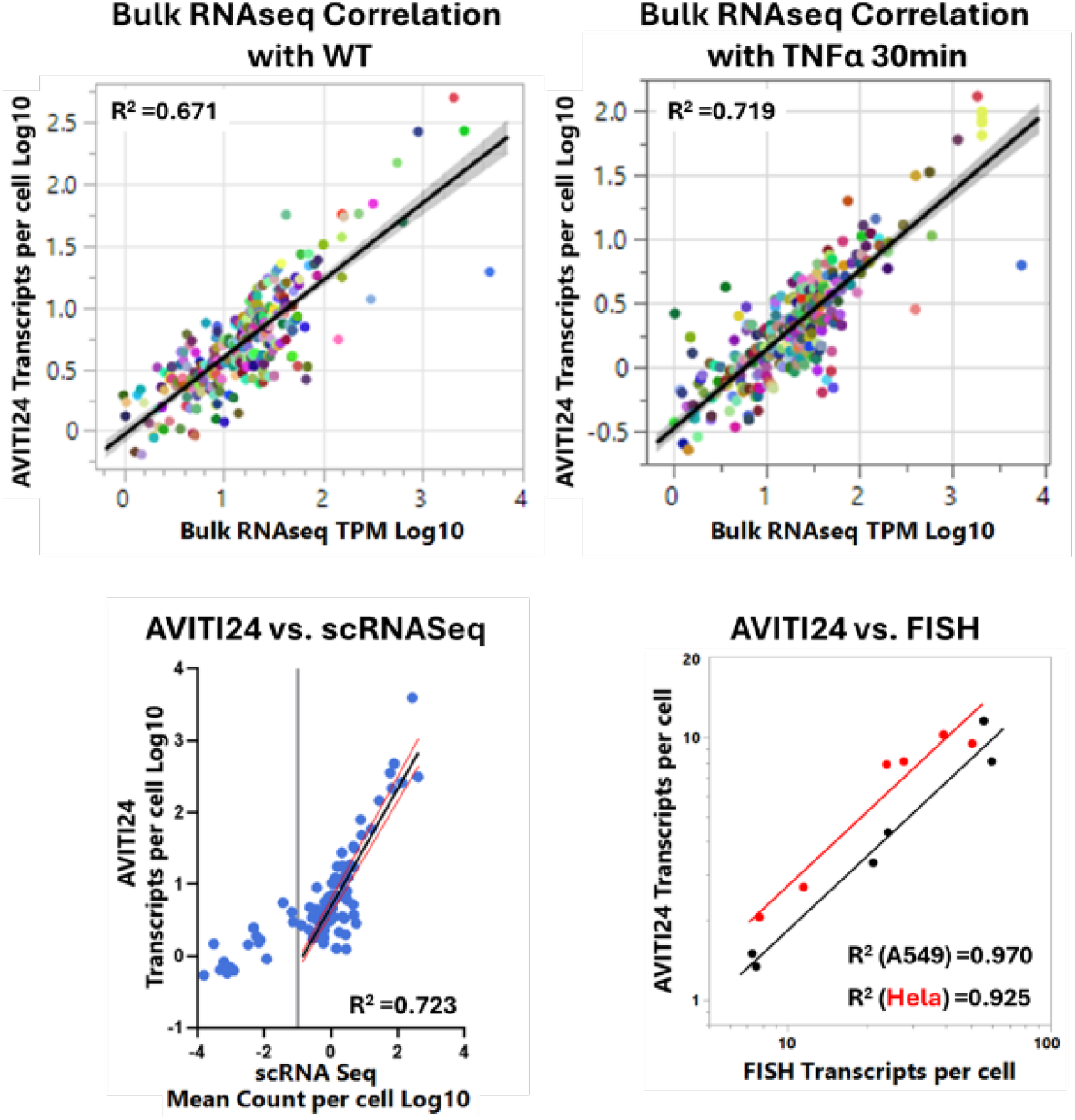
Teton RNA detection on AVITI24 is concordant with RNA-seq, single-cell RNA-seq, and FISH. Teton RNA abundance measurements were validated against three orthogonal methods. (Top left and top right) Comparison with bulk RNA-seq (TPM) across two independent experiments showed strong correlation, with R^2^ values of 0.671 and 0.719, respectively.(Bottom left) Mean gene expression across shared targets was compared between AVITI24 and single-cell RNA-seq (scRNA-seq), yielding an R^2^ of 0.723 for targets above 1 TPM. (Bottom right) RNA transcript counts per cell measured by Teton showed close agreement with RNA FISH in HeLa and A549 cells, with R^2^ values of 0.925 and 0.970, respectively. Together, these results demonstrate that Teton delivers accurate and reproducible RNA quantification across diverse experimental platforms

### Morphological Quantification

To validate the morphological features captured by Teton, we compared cell paint signals to commercially available markers. We assessed localization of the cell membrane, nucleus, actin, mitochondria, endoplasmic reticulum (ER), and Golgi using Teton probes alongside conventional reagents such as wheat germ agglutinin (WGA), RedDot2, phalloidin, Mitotracker, anti-calreticulin, and anti-GM130. Representative images acquired on AVITI24 are shown in Figure 5; comparative results are presented in Supplemental Figure 6. Pixel intensity correlations between the two methods ranged from 0.64 to 0.92, confirming accurate detection of cellular structures by Teton cell paint probes. Segmentation accuracy was further evaluated using expert-annotated images of membrane, nucleus, and actin channels, which serve as the primary input features for cell segmentation onboard the AVITI24 platform. F1 scores, which balance precision and recall, ranged from 0.87 to 0.96, indicating strong agreement with expert-defined boundaries.

**Figure 5:**
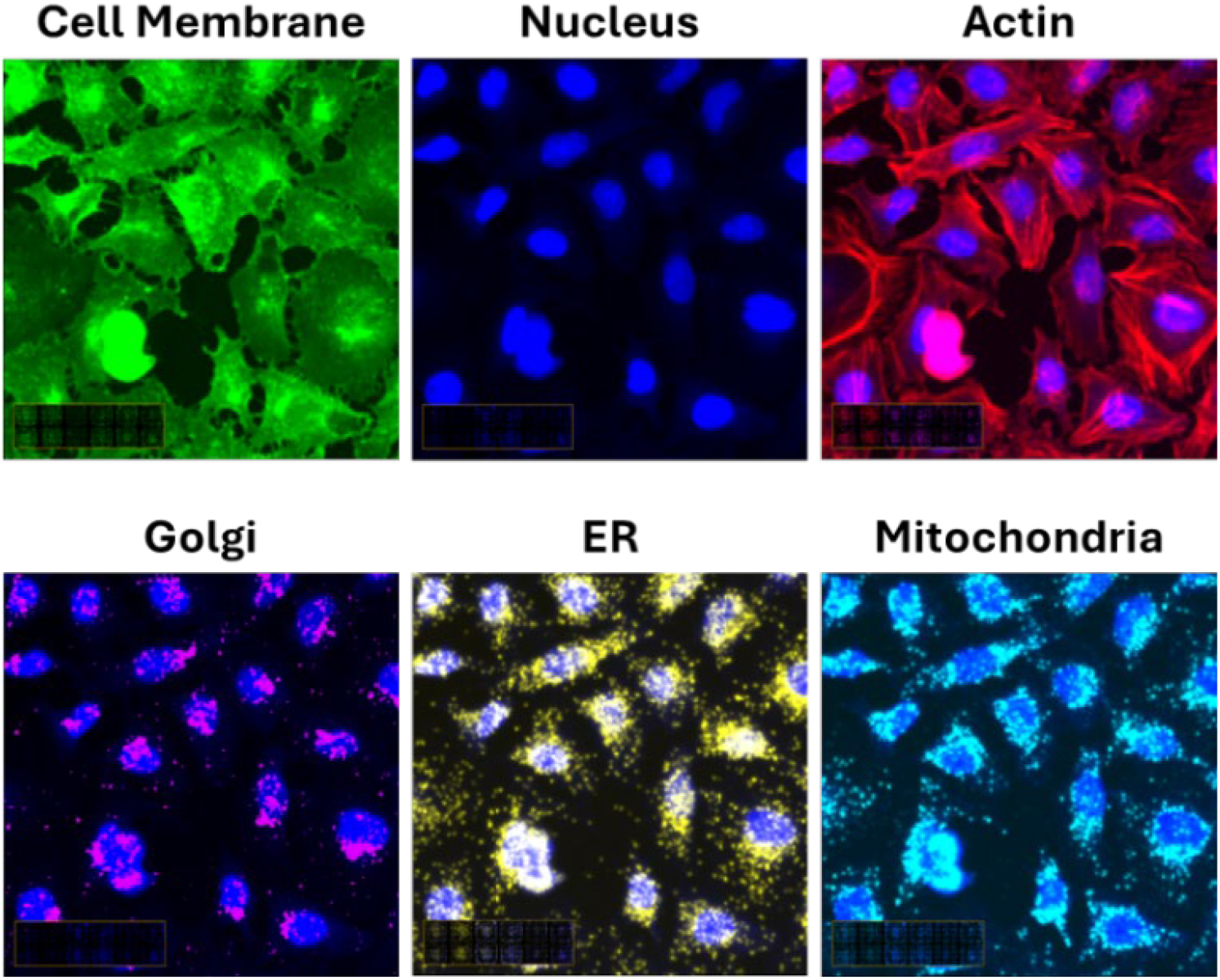
Representative Cell Paint images of HeLa cells acquired using the AVITI24 platform. The six core morphological features include the plasma membrane (green), nucleus (blue), actin cytoskeleton (red), Golgi apparatus (magenta), endoplasmic reticulum (yellow), and mitochondria (cyan). These features enable high-content phenotyping and provide the structural framework for accurate single-cell segmentation and multiomic data integration

### Reproducibility and Robustness

To evaluate the technical reproducibility and phospho-specific detection performance of Teton, we conducted 16 replicate runs across four instruments for the full MAPK/Apoptosis panel (Supplemental Figure 1). Each flowcell was loaded with untreated HeLa cells in one lane and HeLa cells treated with Lambda Protein Phosphatase (LPP) in the other. LPP is a Mn^2^+-dependent enzyme that broadly removes phosphate groups from serine, threonine, and tyrosine residues. This design enabled assessment of both run-to-run reproducibility and the specificity of phosphoprotein signal detection across independent experiments. Reproducibility was high, with an average pairwise R^2^ of 0.975 across all 16 runs and a minimum R^2^ of 0.91 (Supplemental Figure 2). Well-to-well reproducibility was similarly strong, with R^2^ > 0.98 for replicate wells. All runs met Teton quality specifications, including a minimum barcode assignment rate >70%.

To assess phospho-specific signal detection directly, we examined phosphoT197-PKA levels in untreated versus LPP-treated wells across all 16 runs. This site is a well-characterized target of phosphatase activity and served as a benchmark for phospho-signal sensitivity. The mean downregulation of phospho-T197 PKA was −60.1%, with individual values ranging from -54.0% to -68.0%. The coefficient of variation was 7.8%, and the standard deviation across runs was 4.7%, underscoring the platform’s sensitivity, robustness and reproducibility in detecting phosphorylation state changes.

### Teton performance metrics

For the platform to deliver high-throughput, multiomic data, each polony must be accurately linked to its target transcript or protein via barcode sequencing. To validate this capability, we designed an experiment profiling two protein batches and six RNA batches in a 48-well flowcell. The overall barcode assignment rate was 84 %, reflecting a high proportion of successfully decoded polonies. To evaluate false positive rates, barcode identification was repeated with the inclusion of “decoy” barcodes, synthetic sequences absent from the experiment but matched in design to true barcodes. Detection of these decoys provides a proxy for erroneous assignments, which ranged from 2.3 × 10−^6^ to 1.9 ×10 −^4^ in a representative run, confirming high barcode specificity. In total, 357,038,144 polonies passed filtering and were assigned to valid barcodes. At the batch level, assignment rates ranged from 75% to 92%, yielding between 33 million and 57 million assigned polonies per batch. AVITI24’s filtering approach ensures that only confidently assigned, target-specific signals contribute to downstream analysis. To test flowcell capacity, Jurkat suspension cells, which are smaller than HeLa, were seeded into a one-well format. A total of 2,878,435 cells were successfully profiled in 24 hours after fixation, with all platform performance specifications, including assignment rate and mismatch rate, maintained at levels comparable to lower cell density runs.

### A model system to study TNFα Stimulation

To demonstrate the capabilities of our multimodal assay, we selected TNFα stimulation as a model system because of its well-characterized pleiotropic signaling pathways from our panel (Supplemental Figure 1), its role in inflammation and apoptosis, and its relevance to therapeutic intervention. The pathway has been extensively studied across multiple cell types, providing a strong foundation for validating multiomic measurements. In this experiment, HeLa, HepG2, and A549 cells were co-cultured and treated on a 48-well flowcell, yielding over 186,700 total cells, with cell-type density varying based on seeding concentrations (Supplemental Figure 7). After stimulation with TNFα (100 ng/mL) and treatment with kinase inhibitors, the flowcell was assembled and processed on the AVITI24 platform. Barcode-derived target identities were overlaid on Cell Painting images of membrane, actin, and nucleus features that were used for segmentation. This provided a more robust framework than traditional nuclear expansion or two-channel segmentation approaches. UMAP projections from the MAPK/apoptosis panel’s (Supplementary Figure 1) RNA and protein expression profiles revealed clustering by cell type, cell cycle phase, and treatment timepoint, illustrating Teton’s capacity to distinguish both basal states and dynamic responses (Figure 6). This cell stratification was made possible by our single-cell measurements, which preserved the heterogeneity that would otherwise be obscured in bulk analyses where signals are averaged across the populations. Using our multimodal assay, we not only recapitulated established TNFα biology but also revealed cell type specific regulatory differences in pathway response as detailed below.

**Figure 6:**
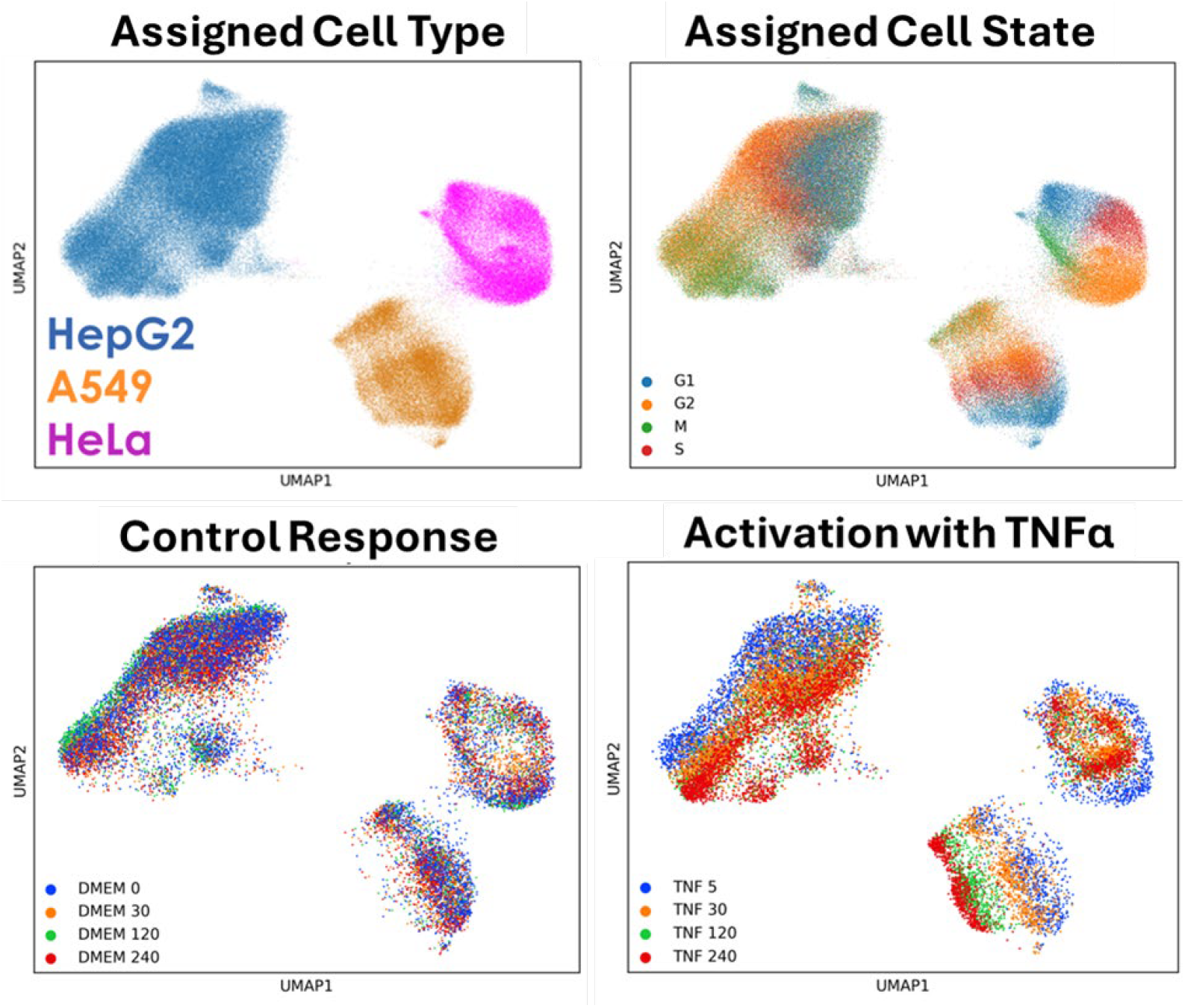
UMAP projections reveal cell type, cell cycle state, and TNFα response dynamics. (Top left) RNA and protein expression profiles were used to assign individual cells to HeLa, A549, or HepG2 clusters. (Top right) Cell cycle states were classified based on RNA expression of canonical G1,

### Phosphorylation Dynamics in Response to TNFα Stimulation

To investigate how TNFα stimulation and kinase inhibition affect signaling dynamics across different cancer cell types, the responses of HeLa, HepG2, and A549 cells within the coculture were profiled across treatments and timepoints. TNFα stimulation induced widespread phosphorylation changes across all three cell types, including upregulation of phospho-T180-p38, phospho-S82-HSP27, phospho-S78-HSP27, and phospho-S46-p53 (Figure 7, Supplemental Figure 8). The well-documented role of HSP27 in stress-response signaling was evident, with p38-driven phosphorylation enhancing its chaperone and anti-apoptotic functions [38,39]. Notably, p38 inhibition by SB202190 (SB) completely suppressed phospho-HSP27, confirming its dependence on p38 activity. In contrast, phospho-p53 and phospho-p38 levels remained largely unaffected by p38 inhibition, suggesting that p38 inhibition selectively targets HSP27-linked pathways without disrupting other TNFα-induced phosphorylation events [40,32]. This model system also served to validate Teton’s ability to measure coordinated changes in protein phosphorylation, RNA expression, and morphology (Figure 7).

**Figure 7:**
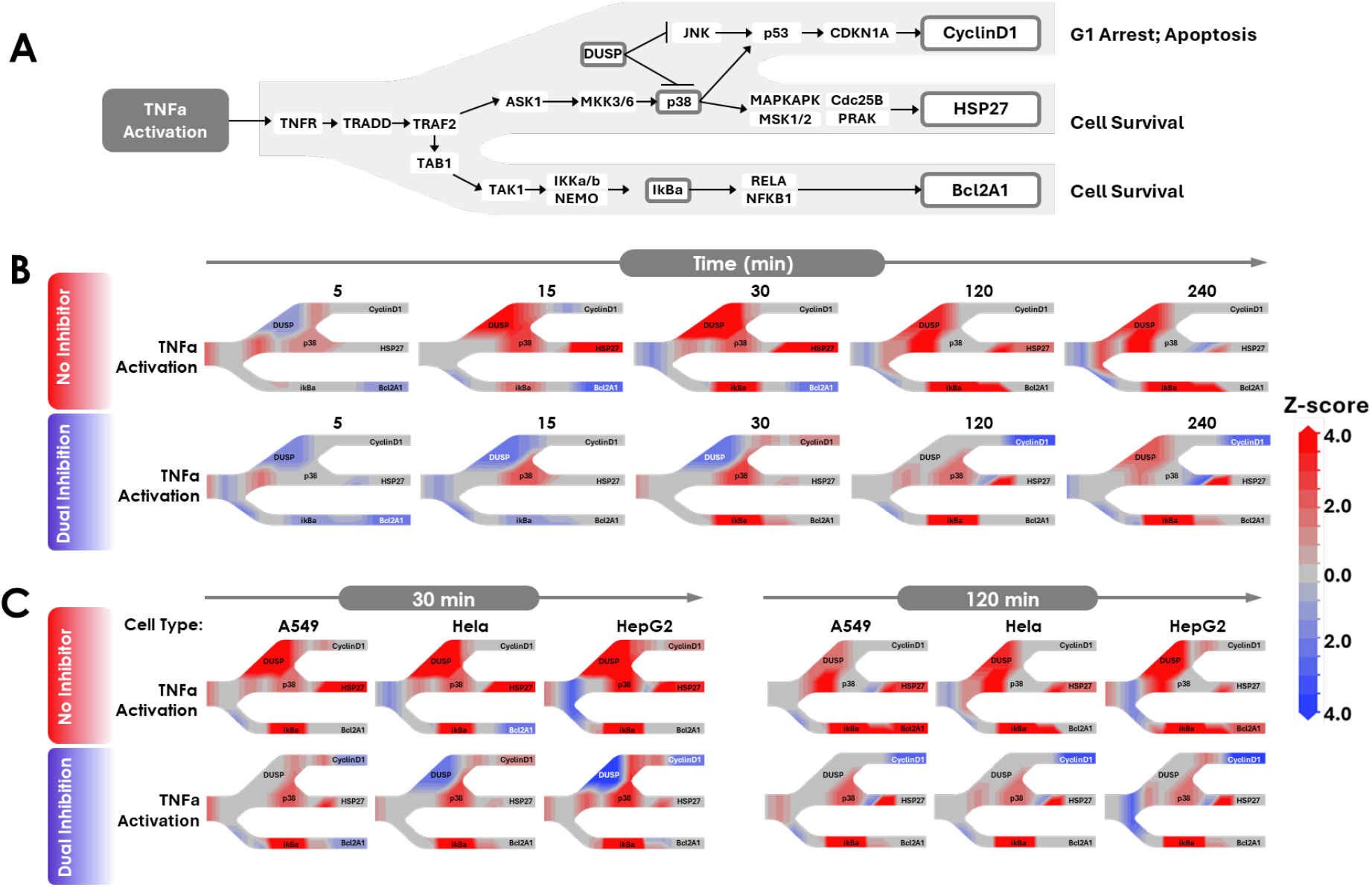
Temporal regulation of TNFα signaling revealed by Teton. (A) Simplified schematic of the TNFα signaling cascade highlighting a curated subset of targets from our MAPK / Apoptosis panel (Supplementary Figure 1) to illustrate key regulatory branches. While Teton profiles hundreds of RNAs, proteins, and phosphoproteins, this view focuses on three dominant pathways: G1 arrest and apoptosis (top), cytoskeletal stabilization via HSP27 (middle), and transcriptional survival signaling via BCL2A1 (bottom). Key mediators, including kinases (ASK1, p38), phosphatases (DUSP family), transcription factors (RELA, p53), and downstream effectors (Cyclin D1, HSP27, BCL2A1), are shown to contextualize dynamic responses. (B) Z-score heatmaps for HeLa cells under TNFα or dual inhibition (SB202190 + FR180204) across representative timepoints (5–240 min). Early timepoints (5–30 min) capture rapid phosphorylation and transcript shifts, such as p38 and phospho-HSP27 induction and early DUSP expression. Later timepoints highlight sustained responses including Cyclin D1 downregulation and modulation of survival pathways. This temporal resolution is critical for capturing pathway sequence and transient inhibitor effects. (C) Comparative heatmaps at 30 and 120 minutes across HeLa, A549, and HepG2 cells with and without dual inhibition. Phospho-HSP27 was strongly induced in HeLa and A549 but modest in HepG2, consistent with its role in cytoprotection [41]. A549 showed marked BCL2A1 upregulation, while HepG2 exhibited the strongest phospho-JNK activation. DUSP expression patterns were cell-type dependent: DUSP2 was preferentially upregulated in HeLa, DUSP6 in HepG2, while DUSP1, DUSP5, and DUSP16 were broadly responsive. Cyclin D1 was consistently downregulated under stress across all models [42]. For all panels, Z-scores reflect the most significantly regulated modality per target (RNA, protein, or phospho-protein), providing a data-type-agnostic view of pathway dynamics. This integrated approach highlights Teton’s ability to resolve complex, time- and cell-type-specific responses to TNFα and kinase inhibition. See Supplemental Figure 10 for a dynamic GIF visualization across all cell types and treatments.

### TNFα-Driven Transcriptional Reprogramming and Apoptotic Signaling

To complement the observed phosphorylation events, we next examined how TNFα stimulation alters gene expression and apoptotic signaling programs across cell types. Robust transcriptional shifts were observed following TNFα activation across all three cell types. The increase in expression of BCL2A1 in Hela and A549 upon TNFα treatment (p = 0.0027 and 0.0050 at 30min, respectively) indicates that cells rely on BCL2A1-mediated survival to evade apoptosis under inflammatory stress (Figure 7) [33]. Upon addition of both SB and FR inhibitors, BCL2A1 expression returned to baseline indicating the pathway was no longer viable for cell survival (Figure 7). In HepG2, pro-apoptotic regulators such as BBC3 (p = 1.0e-11 at 180min in A549) were significantly upregulated, consistent with early activation of the intrinsic apoptotic pathway [43]. Anti-apoptotic targets were suppressed under inhibitor treatments: for example, BCL2 was strongly downregulated in HepG2 (p = 3.2e-06 at 15min with SB), and in A549 and Hela across multiple treatment conditions (P < 6.3e-05), indicating loss of survival buffering [44]. Similarly, CFLAR, an apoptosis checkpoint protein, was downregulated in all three cell types under inhibitor conditions (p = 1.7e-05 to 1.2e-10), consistent with caspase sensitization [45,7].

MCL1 was significantly upregulated in HepG2 upon TNFα treatment (p = 3.5e-05 at 45min), aligning with its reported role in hepatocyte survival under inflammatory stress [8]. In the presence of inhibitors MCL1 remained at baseline. By contrast, it remained largely unchanged in A549. NF-κB negative feedback was supported by the consistent upregulation of NFKBIA, with p = 1.0e-26 in A549, 5.2e-79 in HepG2, and 3.7e-22 in HeLa at 30min post-TNFα treatment (Supplemental Figure 8) [46]. Taken together, the protein, protein phosphorylation, and transcriptional changes detected by Teton upon TNFα treatment are concordant with previously published data.

### Cell-Type-Specific Feedback Regulation of MAPK Targets

The MAPK signaling cascade is one of the most well-studied pathways downstream of TNFα activation, offering a useful benchmark to validate transcriptional and protein-level responses. In HeLa cells, we observed expected MAPK feedback responses, including robust upregulation of dual-specificity phosphatases (DUSPs) and MAPK-responsive transcription factors. DUSP1, DUSP5, and DUSP10 were all strongly induced following TNFα treatment across all three cell types, peaking at 45 minutes (p = 3.2e-05 to 2.1e-41). Inhibitor treatments abolished DUSP1 and DUSP5 induction but did not affect DUSP10, suggesting distinct regulatory control. HeLa also uniquely upregulated DUSP2 (p = 5.2e-13 at 45 min), consistent with cell-type–specific MAPK tuning (Figure 7).

Divergent patterns emerged in other lineages. In A549, DUSP4 was significantly downregulated upon inhibitor treatment (p = 1.9e-08 to 7.6e-16 at 180 min), whereas DUSP6 was selectively upregulated in HepG2 under TNFα stimulation (p = 4.1e-10 at 45 min), consistent with prior reports of its hepatocyte-specific role in ERK1/2 regulation (Figure 7) [47]. These observations underscore the context-dependent nature of MAPK negative feedback.

MAPK-responsive transcription factors followed both expected and cell-specific expression trajectories. FOS was strongly upregulated in all three cell types at 45 minutes (p = 1.7e-30 to 3.6e-83), reflecting its role as a canonical immediate-early gene. FR treatment nearly abolished FOS induction (p = 0.38 in A549, 0.68 in HepG2, 0.0015 in HeLa), while SB had a partial effect. Notably, HepG2 showed relatively weak TNFα-driven FOS induction overall (p = 0.069), suggesting enhanced sensitivity to p38 inhibition in this context. JUN was consistently upregulated across cell types and timepoints (P < 0.045), indicating sustained MAPK engagement [48,49].

GADD45 family members also showed lineage-specific regulation. GADD45B, a known JNK activator, was selectively upregulated in HepG2 after TNFα stimulation (p = 1.0e-16 at 120 min) [50,51], while in A549 and HeLa, GADD45B expression was mainly induced under SB inhibition, implying that p38 suppression unmasks compensatory JNK activation. GADD45A followed a similar pattern: strongly induced in HeLa and HepG2 with TNFα alone (p = 4.2e-06 and 5.2e-09) but only upregulated in A549 under inhibitor conditions.

Together, these findings illustrate that Teton reliably detects known MAPK signaling responses in well-characterized systems like HeLa, while also resolving cell-type–specific transcriptional adaptations in A549 and HepG2.

### Kinase Inhibition Disrupts Proliferation and MAPK Feedback

To investigate how kinase inhibition alters MAPK feedback and proliferative signaling, we examined the transcriptional and proteomic consequences of SB202190 (SB) and FR180204 (FR) treatment across cell types. These conditions revealed coordinated downregulation of proliferative drivers and MAPK-associated scaffolds, suggesting a transition toward apoptosis. MYC, a critical transcriptional regulator of growth, was sharply suppressed in HepG2 following p38 inhibition (p = 4.4e-11 at 45min with SB), consistent with p38’s reported role in stabilizing MYC under stress conditions [52]. This was accompanied by significant protein-level suppression of CyclinD1 in all three lineages with TNFα + inhibitors (p = 0.033 in A549, 0.0035 in HeLa, 4.2e-06 in HepG2 at 180min), aligning with reduced cell cycle progression (Figure 7). Further support for cell cycle arrest came from phospho-CDK1, which was strongly suppressed at the protein level in A549 (p = 2.5e-07) and HepG2 (p = 0.010) as early as 10min following TNFSB treatment, indicating G1/S checkpoint blockade [53].

At the transcriptional level, p38 inhibition also disrupted signaling crosstalk with the JNK pathway. GADD45B, a stress-responsive gene known to activate JNK, was selectively upregulated in HepG2 upon TNFα stimulation (p = 1.0e-16 at 120min), consistent with models where p38 sustains JNK signaling via GADD45B [28]. However, this axis appeared dismantled under inhibitor treatment, as both MAPK8IP3 and MAPK8IP1, scaffold proteins critical for JNK assembly were sharply downregulated in A549 and HepG2, respectively (p = 3.4e-08 and 3.8e-23 at 90min), consistent with prior reports [54] and confirmed by our data.

Broader transcriptional remodeling was also evident following dual ERK1/2 and p38 inhibition. The pro-apoptotic gene BAD was significantly suppressed in HepG2 under TNFFR+SB (p = 4.8e-04 at 90min), suggesting loss of mitochondrial priming [28]. BIRC5 (Survivin), a key mitotic regulator, was consistently downregulated in A549 (p = 6.9e-05), HepG2 (p = 5.8e-04), and HeLa (p = 0.025 at 45min), indicating compromised mitotic progression. MAPK13, a noncanonical p38 isoform, was also suppressed in A549 (p = 0.022) and HeLa (p = 1.9e-04 at 60min), underscoring auxiliary MAPK suppression. Notably, phosphatase feedback loops also collapsed under combined inhibitor pressure: transcripts for DUSP2, DUSP6, and DUSP9 were coordinately suppressed, reflecting a broader loss of regulatory buffering within the MAPK network.

### Cell Paint Responses

Cell Paint features enabled robust delineation of cell types and treatment responses based on intracellular morphology. Morphological distinctions were also evident within cell types across cell cycle phases. In A549, average area decreased from G1 (4,896) to M phase (4,122), while HeLa displayed a similar trend from G1 (5,446) to M (3,621). HepG2 exhibited less fluctuation but trended smaller overall, with G1 cells averaging 3,563 and M phase cells 3,401. These measurements suggest that cell area correlates with both cell type and state, reinforcing the utility of Cell Paint for morphological profiling.

Dual inhibition led to marked reductions in ER signal across all three cell types: from 760,364 to 484,739 in A549, 1.05 × 10^6^ to 669,075 in HeLa, and 885,321 to 635,737 in HepG2, consistent with ER destabilization upon MAPK pathway suppression. Golgi signal was also disrupted under inhibitor treatment, decreasing from 279,373 to 204,853 in A549, 364,189 to 275,068 in HeLa, and 268,645 to 198,680 in HepG2. These observations align with previous studies showing that MAPK signaling regulates ER and Golgi integrity, and that Golgi-localized proteotoxic stress can trigger fragmentation and transcriptional remodeling [55,56]. Cell area decreased following inhibitor treatment in nearly all cell types and cell cycle states, except for HepG2 in G2 and M phase. No significant changes were observed in any of the six morphological features in DMEM-treated control wells, confirming that the observed phenotypes were treatment-specific.

## Conclusion

In this study, we employed TNFα stimulation as a model system to achieve two objectives: first, to confirm that the Teton assay on AVITI24 generates data consistent with established biological responses; and second, to demonstrate its unique ability to reveal complex, cell-type-specific regulatory mechanisms underlying broader signaling networks. Teton simultaneously profiles cellular morphology, RNA expression, protein expression, and phosphoprotein dynamics in the same cells without the need for library preparation. It integrates a compartmentalized flowcell for cell culture and perturbation with a cell imaging and sequencing instrument, enabling completion of all assays within 24 hours. The platform supports morphological profiling of up to six subcellular components, concurrent measurement of approximately 350 RNA targets and 200 proteins, including phosphoproteins, across millions of cells, and up to 48 experimental conditions per run.

Extensive validation against orthogonal technologies, including immunofluorescence, western blotting, RNA-seq, scRNA-seq, and FISH, confirmed Teton’s accuracy. These comparisons indicate that the data generated reflects genuine biological responses.

We used the platform to examine TNFα-induced signaling pathways and their modulation by kinase activity, yielding detailed insights into molecular dynamics. Although each modality was analyzed independently in this study, the high degree of cross-modal agreement reinforces the biological conclusions. Our results revealed p38-driven phosphorylation as critical for modulating stress responses, notably demonstrated by the complete suppression of phospho-HSP27 following p38 inhibition. Additionally, kinase inhibition significantly affected apoptotic signaling and cell cycle regulation: pro-apoptotic factors like BBC3 were upregulated, anti-apoptotic regulators including BCL2 and CFLAR were downregulated, and key cell-cycle mediators (MYC, Cyclin D1, phospho-CDK1) were suppressed, reinforcing p38’s role in stress-mediated cell cycle arrest.

We also observed cell-type-specific expression patterns, including differential regulation of DUSP phosphatases and MAPK-responsive transcription factors such as FOS. Our data further identified significant crosstalk between p38 and JNK signaling pathways and demonstrated that dual inhibition of p38 and ERK1/2 disrupted multiple cellular programs. These findings elucidate mechanisms by which cancer cells evade TNFα-induced apoptosis not by failing to detect TNFα but by kinase-dependent rewiring of intracellular signaling that buffers stress and apoptotic signals.

Teton on AVITI24 provides a framework for generating detailed, temporally resolved pathway maps that can deepen mechanistic understanding of cellular biology and perturbation responses, with potential applications in targeted drug discovery. Future efforts will focus on integrating the multimodal data as input for foundation model training to extract additional biological insights from these rich datasets.

## Methods

### Cell Culture and Treatment

HeLa (ATCC, CCL-2), A549 (ATCC, CCL-185), and HepG2 (ATCC, HB-8065) cell lines were obtained from the American Type Culture Collection (ATCC) and maintained under standard conditions. HeLa cells were cultured in Dulbecco’s Modified Eagle Medium (DMEM; ThermoFisher, 10569010) supplemented with 10% heat-inactivated fetal bovine serum (FBS; ThermoFisher, A3840202), 100 U/mL penicillin-streptomycin (ThermoFisher, 15140122), 1 mM sodium pyruvate (ThermoFisher, 11360070), and 1× MEM non-essential amino acids (ThermoFisher, 11140050). A549 cells were cultured in F-12K Nutrient Mixture (Corning, 10-025-CV), and HepG2 cells in Eagle’s Minimum Essential Medium (EMEM; Quality Biological, 112-018-101), with both media formulations supplemented with 10% FBS and 100 U/mL penicillin-streptomycin. For multi-lineage coculture experiments, cells were combined in a 25% HeLa, 25% A549, and 50% HepG2 ratio and seeded into custom 48-well flowcells at a total density of ∼4000 cells per well. After 16-20 hours of incubation at 37°C and 5% CO_2_, treatments were introduced using fresh DMEM containing 100 ng/mL TNFα (Stemcell Technologies, 78068), with or without 100 μM SB202190 (p38 inhibitor; Sigma, S7067-5MG) and 100 μM FR180204 (ERK inhibitor; Sigma, SML0320-5MG). Treatments were staggered across timepoints (0 to 240 minutes) to enable synchronized assay completion. Following stimulation, cells were fixed in 3.7% formaldehyde for 30 minutes at room temperature and stored at 4°C in 0.1 U/μL RiboLock RNase Inhibitor (ThermoFisher, EO0384) for downstream processing.

### Bulk RNA Sequencing

For bulk RNAseq, live cells were washed twice with 150 µL of 1X PBS. The PBS was removed, and 200uL of Tri reagent (Zymo Research, R2050-1-50) was added to lyse the cells. After agitation via pipette, RNA was extracted from the lysed cell suspension by following manufacturer’s protocol for the Direct-zol RNA Miniprep kit (Zymo Research, R2052). Following isolation of RNA, 50ng of total RNA for each sample was used for RNA-seq library prep with the Watchmaker RNA library prep kit with Polaris Depletion (7BK0002) per manufacturer’s protocol. Briefly, total RNA was depleted of rRNA and then fragmented at 85°C for 5 minutes. The fragmented RNA was reverse transcribed and A-tailed. Adapters with unique indices were ligated onto each sample and then amplified by PCR. The libraries were then pooled and sequenced on AVITI (Element Biosciences).

### Single Cell RNA Sequencing

Live cells for single cell analysis were obtained by trypsinizing adherent cells to form a cell suspension. Cell media was exchanged to PBS by centrifuging (400xg 5min), decanting, and resuspending in 1x PBS. Cells were filtered through a 40um cell strainer (Fisher, 07-001-106) to remove debris and aggregates, and were counted via hemocytometer. The final cell suspension was prepared in PBS with 0.04% BSA at a final concentration of 1000 cells/ uL. Single-cell suspensions were processed for single-cell RNA sequencing using the 10x Genomics Chromium 3’ reagent kit (v3). Single-cells were collected into individual droplets using the 10x Genomics Chromium Controller. Following droplet generation, reverse transcription (RT) was performed within each droplet to add cell and molecule-specific barcodes. The cDNA was subsequently fragmented, ligated with adapters for sample indexing, and then amplified using a PCR-based approach to generate libraries for each sample. Samples were pooled and then sequencing was performed on AVITI (Element Biosciences).

### Fluorescence in situ hybridization (FISH)

For FISH detection, Hela, HUVEC and A549 cells were fixed with 4% formaldehyde for 30 minutes at room temperature. Fixed cells were washed three times with 150 µL of 1X PBS and permeabilized with 100 µL of 70% ethanol for 10 minutes at room temperature. FISH probes were designed and ordered using Stellaris custom probe designer (Biosearch technologies). FISH probes were hybridized and imaged according to manufacturer’s recommendations. Probe targets included NRAS, KRAS, TP53, PIK3CA, EGFR, and BRCA2.

### siRNA Knockdown

For siRNA-mediated gene knockdown, HeLa cells were seeded one day prior to transfection and allowed to reach 60-70% confluence. On the day of transfection, Opti-MEM medium (ThermoFisher, 31985070) was prewarmed to 37°C. Lipid-siRNA complexes were prepared in Opti-MEM, with each siRNA transfection performed individually. Briefly, two separate 0.6 mL Eppendorf tubes were prepared: one containing 22.5 µL of warm Opti-MEM mixed with 0.9 µL of 10 µM siRNA (9 pmol total), and the other containing 22.5 µL of warm Opti-MEM mixed with 2.7 µL of Lipofectamine RNAiMax Reagent (ThermoFisher, 13778030). The two solutions were gently combined and incubated for 5 minutes. For lipid-only control wells, the preparation followed the same steps, excluding the siRNA. Cells were washed twice with 1X PBS, and 85 µL of Opti-MEM was added to each well. Next, 15 µL of the lipid-siRNA complex (or lipid-only control) was added per well, and the plate was gently mixed. After four hours of incubation, the media was replaced with DMEM containing 10% FBS to support cell growth. Untreated control wells also underwent media changes at this step. Cells were maintained for 72-96 hours before fixation for downstream assays.

### Immunofluorescence Validation

For immunofluorescence validation, cells were fixed with 3.7% formaldehyde for 30 minutes at room temperature and stored at 4°C. Fixed cells were washed once with 150 µL of 1X PBS and permeabilized with 100 µL of 70% ethanol for 10 minutes at room temperature without pipetting. Following three PBS washes, primary antibodies were diluted according to the manufacturer’s recommendations in 1X PBS, and 50 µL of the diluted antibody solution was added to each well. The cells were incubated at room temperature for one hour, followed by three PBS washes with a five-minute interval between each. A fluorescent secondary antibody (2.5 µg/mL) was then applied for one hour at room temperature. After three additional PBS washes, cells were stored in 100 µL of PBS and imaged using fluorescence microscopy.

### Western Blot

Cells were passaged into T75 flasks at a density of 1 million cells for overnight growth. Prior to treatment, fresh media containing the appropriate dilution of treatment factors was prepared. Old media was aspirated, and treatment media was added to cover the cell monolayer, followed by incubation at 37°C for the designated time. After treatment, media was removed, and cells were washed with 0.25% trypsin, incubated until detachment, and then quenched with media before transfer to a conical tube. A 10 µL aliquot of the cell suspension was taken for counting using a hemocytometer. Cells were pelleted by centrifugation at 200 rcf for 5 minutes, media was aspirated, and pellets were resuspended in Trizol reagent (Zymo Research, R2050-1-200) at an appropriate volume for downstream processing or storage at -80°C.

For protein extraction, lysates were mixed with an equal volume of 200-proof ethanol, followed by the addition of four volumes of pre-cooled acetone. Samples were incubated on ice for 30 minutes and centrifuged at 1000 x g for 10 minutes. The supernatant was discarded, and the protein pellet was washed with 200-proof ethanol, centrifuged again, and air-dried for 10 minutes before resuspension in PBS. If resuspension was difficult, sonication was used. Samples were stored at 4°C for immediate use or -20°C for future experiments.

For Western blotting, lysates were treated with a 1X reducing agent and heated at 90°C for 10 minutes. Proteins were separated using a 4-12% Bis-Tris NuPAGE gel (ThermoFisher, NP0322BOX) at 180V for 40 minutes. Following electrophoresis, transfer was performed using the iBlot device under P0 conditions (20V, 7 minutes) for proteins 30-150 kDa, with extended transfer times for larger proteins. Membranes were briefly rinsed in DI water, cut as needed, and blocked in 5% skim milk for 15 minutes at room temperature. After blocking, membranes were washed in TBST for 15 minutes and incubated with the primary antibody solution for 1 hour at room temperature. Following incubation, membranes were washed in TBST for 15 minutes, then incubated in secondary-HRP antibody diluted 1:5000 in 5% skim milk for 1 hour at room temperature. Membranes underwent four sequential TBST washes (15 minutes, 5 minutes, 5 minutes, 5 minutes) before a 5-minute incubation in substrate mix (1:1 ratio of substrate and enhancer). Chemiluminescence imaging was performed immediately using a Typhoon imager. To reduce background, membranes were washed in TBST before imaging. For Typhoon imaging, the stage was cleaned, and the filter module was switched to “Chemilum” by removing the first filter. In the fluorescence menu, both Cy2 and chemiluminescence methods were selected. Cy2 settings included a PMT of 500V, a pixel size of 50 µm (or 100 µm for strong bands or large gels). Primary antibodies used included: anti-TFEB [EPR22941-6] (Abcam, ab267351), anti-Cdk4 [EPR4513-32-7] (Abcam, ab108357), anti-CD44 [RM1084] (Abcam, ab316123), anti-beta III Tubulin [EPR19591] (Abcam, ab215037), anti-PMCA1 [EPR12029] (Abcam, ab190355), anti-ICAM1 [EPR24639-3] (Abcam, ab282575), and anti-Hsp27 (phospho S78) [Y175] (Abcam, ab32501).

### Mass Spectrometry

To validate Teton-derived protein expression data, we performed label-free quantitative proteomics on cell lysates prepared from treated and untreated samples. Cells were cultured under standard conditions and subjected to the appropriate treatments. Adherent cells were detached using trypsin, pelleted by centrifugation, and washed twice with pre-chilled PBS to remove residual media. After a final centrifugation step, cell counts were performed to ensure a minimum of 5 million cells per sample. Pellets were transferred to sterile Eppendorf tubes, sealed with parafilm, and stored at –80 °C. For shipment, samples were transported on dry ice to MtoZ Biolabs for downstream analysis.

Protein extraction and digestion were performed using a standardized SP3 protocol. Briefly, proteins were lysed in 8 M urea with PMSF, reduced with dithiothreitol (DTT), alkylated with iodoacetamide (IAM), and neutralized with DTT. Proteins were purified via bead-based capture and digested overnight with sequencing-grade trypsin (Promega) at 37 °C. Peptides were desalted using C18 columns, vacuum-dried, and resuspended in 0.1% formic acid for LC-MS analysis.

Peptide separation was performed on a Thermo Vanquish Neo UHPLC system equipped with an Acclaim PepMap RPLC C18 column (150 μm × 170 mm, 1.9 μm, 100 Å), using a 66-minute gradient from 4% to 95% mobile phase B (80% acetonitrile + 0.1% formic acid) at a flow rate of 600 nL/min. Mass spectrometry was conducted on a Thermo Orbitrap Exploris 480 in positive ion mode. Full MS scans were acquired from m/z 400–1200 at 60,000 resolution, with an AGC target of 300% and a 50 ms maximum injection time. Data-dependent MS/MS fragmentation was performed using higher-energy collision dissociation (HCD) at 30% normalized collision energy, acquired at 15,000 resolution.

Raw data were searched against the Homo sapiens UniProt database with trypsin specificity (up to two missed cleavages), carbamidomethylation (C) as a fixed modification, and oxidation (M) and N-terminal acetylation as variable modifications. Precursor and fragment mass tolerances were set to 20 ppm. Label-free quantification was performed using peptide peak areas, with normalization by the median method and imputation of missing values using half-minimum estimation.

### Probe Formulation

For each mRNA target, specific single-stranded DNA oligonucleotides were designed and annealed to circular DNA molecules with unique barcodes. Similarly, a coordinating molecule specific to rabbit primary antibodies was conjugated to uniquely barcoded circular DNA molecules for protein detection. Pooled RNA (350-plex) and protein (200-plex) probe panels were prepared and either used immediately or stored at -20°C for later use.

For custom protein probe assembly, primary antibodies were diluted to 3.4 μg/mL using the provided Teton Custom Add-On Protein Buffer. After thawing, Teton detection probes were transferred to a new 96-well plate, followed by the addition of the diluted primary antibodies. The mixture was incubated at room temperature for one hour before pooling. The final pooled protein panel was adjusted to a total volume of 1620 μL using Teton Custom Add-On Protein Buffer. The completed protein panel was aliquoted into 200 μL portions and stored at -20°C. Before use, an aliquot was thawed, mixed, and added to the fixed panel protein tube for integration into the spatial multiomics workflow.

### Spatial Multiomics Workflow

The spatial multiomics workflow was performed on the AVITI24 platform. Barcoded protein and RNA probes were sequentially injected into the flowcell, allowing specific binding to their targets in situ. Following probe binding, excess probes were washed away, and the bound barcodes were amplified via rolling circle amplification (RCA). Amplified signals were detected using avidite-based chemistry and decoded. Cell boundaries were delineated using cyto-profiling probes, which enabled accurate mapping of barcode signals to individual cells.

### Cell Paint Segmentation and Morphology Analysis

For each cell paint feature, a Z stack of seven images with 1 um spacing was captured for each field of view. A two-dimensional projection was produced by summing the intensities of each pixel across the Z stack. The projections of the cell membrane, nuclear, and actin cell paint features were used to perform cell segmentation. Custom cell segmentation models were trained on Teton cell paint images acquired on AVITI24. A Cellpose architecture was utilized, and models were retrained using manually annotated AVITI24 images [57]. Cell segmentation was performed using nuclear, cell membrane, and actin cell paint images as model inputs and nuclear segmentation was performed using only nuclear cell paint images. Both cell and nuclear segmentation models were trained using a diverse set of images from different cell types. The cell segmentation model was trained using 1,441 images and tested on 315 608x608 images. The nuclear segmentation model was fine-tuned from the Cellpose ‘nuclei’ model using 937 images and tested on 165 images. Following training of the cell segmentation model on all cell types, a specific model was then fine-tuned on HeLa cell images only and was used for segmentation here. The AreaShape module of CellProfiler was used to calculate cell morphology features on the resulting cell segmentation boundaries. Prior to further morphological analysis, the projection for each cell paint feature was registered to the cell membrane projection. Per-cell intensity was calculated for each cell paint feature by first subtracting background and summing the intensity per pixel across the full cell area. Additionally, the MeasureObjectSizeShape, MeasureGranularity, MeasureObjectIntensity, MeasureObjectIntensityDistribution, and MeasureTexture Cellprofiler modules were all executed for determination of additional morphological features [58]. This resulted in a total of 575 morphological features calculated per cell across all six cell paint features and segmentation mask.

### Sequencing Primary Analysis

To limit optical crowding, barcodes were processed in multiple batches, each delineated with a unique sequencing primer. For each barcode sequencing cycle and field of view, a Z stack of 7 images with 1 um spacing was captured for each channel corresponding to the four Avidite dyes. Prior to any further processing for polony location determination or base calling, each Z stack of images was first jointly processed with a CNN model to sharpen features, reduce noise, and remove background. For each slice of the Z stack, an analysis pipeline was used to determine a set of polony locations, as described previously, [37]. After determination of polony locations in each slice, duplicate spots across the Z dimension were resolved based on distance, sequencing identity, and quality. With a set of unique locations in each slice, base calling proceeded as described previously [37]. To summarize, following image-denoising, each sample image was registered to the set of polony locations based on a reference image associated with those locations. An intensity was extracted for each polony location across all Avidite channels. Extracted intensities were corrected for spectral cross-talk and intensities were normalized across channels. A base call for each polony was determined from the maximum normalized intensity per cycle. Following base calling across all sequencing cycles for a given batch, polonies were assigned to the expected barcodes allowing up to two mismatches. To associate assigned barcodes to cells, the set of polony locations for each batch was registered to the cell segmentation result. This was accomplished indirectly by registering a reference image associated with the set of polony locations to the projection image of the cell membrane. The registered polonies were then assigned to individual cells and annotated as either nuclear or cytoplasmic.

### Data analysis

Standard pre-processing of counts data consisted of filtering cells and then normalizing counts. Cells were filtered based on a variety of criteria to remove outliers and eliminate erroneously segmented objects. First, cells were filtered based on area, keeping only cells between the 2.5^th^ and 97.5^th^ percentiles of the cell area distribution. Next, cells were filtered based on total assigned counts in each batch, keeping only cells between the 2.5^th^ and 97.5^th^ percentile of total assigned counts in all batches. Finally, cells with an assignment rate (assigned polonies / total polonies) less than 50% were removed. Cells were filtered independently in each well to account for sample-specific differences in distributions. Within each batch, counts were normalized relative to the total number of assigned counts in each cell. For multi-sample analyses, counts were further normalized between samples. Within each batch, the counts were linearly scaled such that the median ratio to an average pseudo-reference sample was one (*i.e.*, the median of ratios method, [59]). Implementations of filtering and normalization methods are available at https://github.com/Elembio/cytoprofiling.

For differential expression analyses, filtered and normalized counts were averaged within each condition to produce an average count per cell for each target in each condition. For each pairwise comparison, the average count per cell was compared across conditions to produce a log ratio for each target. Under the assumption that most targets involved in the comparison were invariant, the standard deviation of the null distribution of log ratios was estimated as 1.48 times the median absolute deviation. A differential z score for each target was calculated as the log ratio divided by the standard deviation. Then converted to p-values for reporting using a 2 tailed test for a standard normal distribution.

UMAP analysis was performed with the scanpy suite of tools. Per-cell feature counts were log-transformed and then standardized to unit variance and zero mean. Principle components analysis was performed on the scaled features, with the final number of principle components utilized determined from a manual inspection of the variance explained by each PCA axis. Finally, the nearest neighbor distance matrix was computed and used to determine the UMAP projection.

To classify the HepG2, HeLa and A549 cell type mixture, the combined sample was first clustered using the unsupervised Leiden method. When these clusters were visualized on a UMAP projection, three well-separate clusters were observed. To assign and confirm the association of each cluster with a unique cell type, cell paint images from multiple examples of each cluster were manually inspected.

### Classification of Cell Cycle State

Furthermore, in order to leverage the single-cell resolution of Teton and separate cells into their distinct cell cycle state, a classifier was trained using a combination of cell synchronization experiments. Hela cells (ATCC, CCL-2) were seeded onto PLL coated Teton™ CytoProfiling 12 well flow cell at 4500 cells per well in culture media (DMEM (Gibco, 10566-016) plus 10% FBS (Cytiva, SH30396.03HI), 1% PenStrep (Cytiva, SV30010), 1% MEM NEAAs (Gibco, 11140,050), and 1% Sodium Pyruvate (Gibco, 11360-070)) and incubated overnight to allow cells to attach to surface. Media was then replaced with media with cell phase arresting agents for: S (Thymidine, 2mM, Sigma T1895-1G), G2 (RO-3306, 10uM, Sigma 217699-5mg), M (Nocodazole, 0.2ug/ml, Stemcell Technologies 74072), G1 (Lovastatin, 20uM, Sigma 438185-25mg), G1/S (Aphidicolin, 5ug/ml Sigma 178273-1mg) and allowed to incubate at 37°C for 20 hours before fixation. Cells were washed twice with 200uL per well of warmed DPBS and fixed in 150uL per well of 3.7% formaldehyde (Sigma-Aldrich, F8775-25mL) for 20min, then washed twice with 200uL per well 1x PBS then stored at 4°C in PBS containing 0.1U/uL RiboLock RNase Inhibitor (Thermo Scientific EO0381) until used on AVITI24. Flowcell was run on AVITI24 using the Teton Human MAPK & Cell Cycle Kit (860-00023). Two replicate flowcells were run on AVITI24.

Following run completion, transcript counts for both AVITI24 runs were normalized per cell by total detected counts per batch and outlier cells were excluded via cell area and assigned counts as described above. Cells were then combined across both runs from all wells. Further preprocessing normalized cells across all sequencing batches and log-transformed counts. Transcript features were projected into their principal components. Upon inspection of the first two principal components, a two-dimensional annulus was observed, consistent with prior observation (Supplemental Figure 9) [60]. Cells from wells with different synchronization agent treatments were observed to cluster in different regions of the annulus. Unsupervised clustering was performed on the PCAs using the Leiden algorithm to achieve the expected number of clusters for each cell cycle state [61]. Following clustering, clusters were assigned to each cell cycle state based on their position on the annulus and their enrichment of cells synchronized in a specific state. This resulted in four clusters for each cell cycle state. Then an enrichment analysis was performed for each state relative to the others using a t-test method. This resulted in an enrichment score for each transcript in each cluster relative to the others. The top 20 transcripts for each state were then used in other AVITI24 runs to score cells for the most likely cell cycle state via methods described previously [62].

