## Supplementary Figures and Tables for "High-Throughput Multiomics Profiling of Model Systems Using the AVITI24 Platform"

### Supplemental Figures

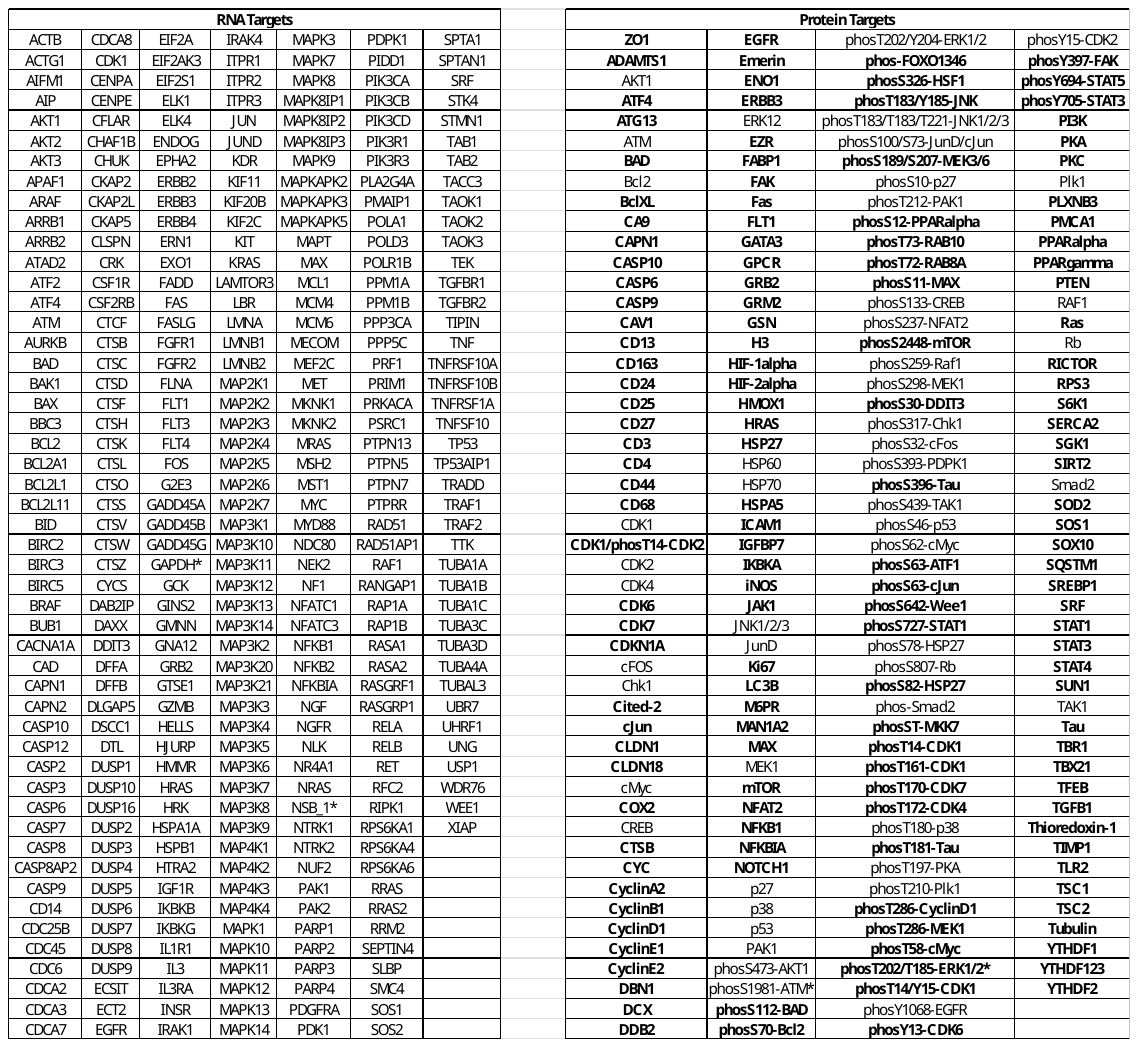

**Supplemental Figure 1:** **Targets included in the high-plex TNFα model system run.**

The AVITI24 Teton MAPK and apoptosis RNA/protein panels (MAPK/APOPRNA) were supplemented with an additional 150 protein targets (bolded) to create the full panel used in this study. Targets marked with an asterisk (*) indicate multiple probes were designed against the same target.

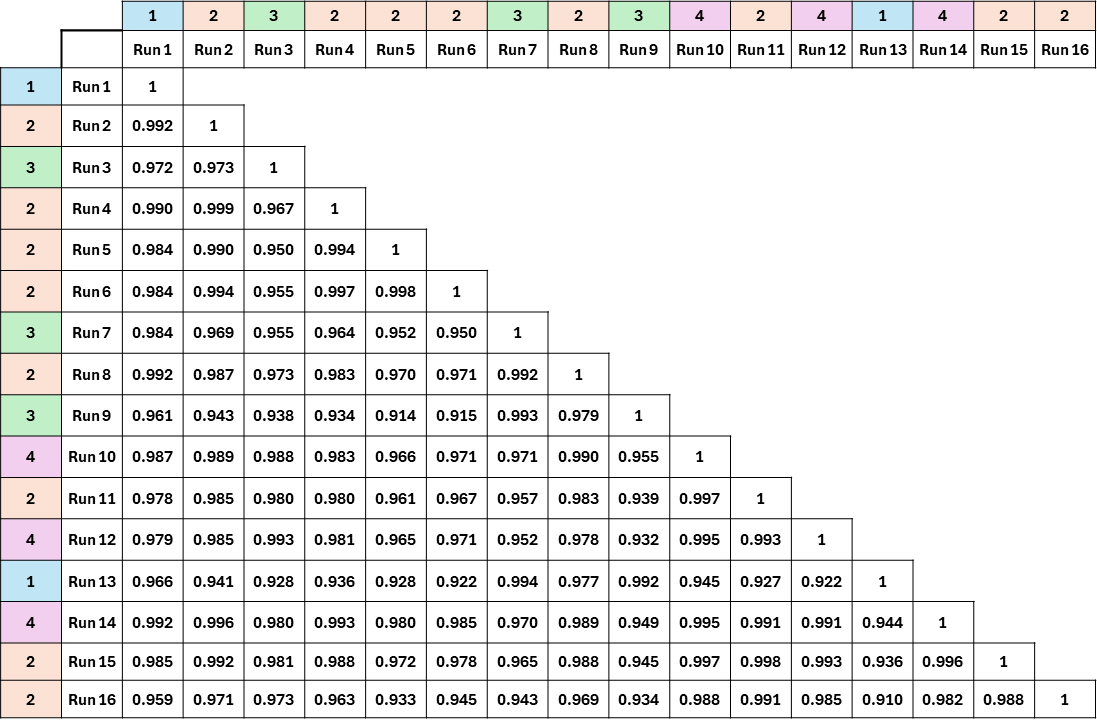

**Supplemental Figure 2: Run-to-Run Reproducibility Across AVITI24 Systems**

Each cell in the matrix shows the R² value for raw counts between a pair of AVITI24 runs. A total of 16 independent runs were compared, spanning four separate AVITI24 instruments. The analysis revealed a high degree of technical reproducibility, with an average R² of 0.975 and values ranging from 0.910 to 0.999. Colored annotations indicate the instrument used for each run, confirming consistency both within and across instruments.

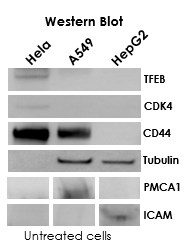

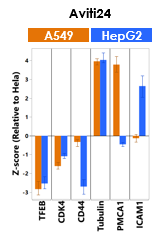

**Supplemental Figure 3: Concordance between Teton and western blot measurements.**

Left: western blot of untreated HeLa, A549, and HepG2 lysates for six targets: TFEB, CDK4, CD44, Tubulin, PMCA1, and ICAM1. Right: Teton-derived Z-scores for A549 and HepG2 cells relative to HeLa, showing strong agreement with western blot signal patterns.

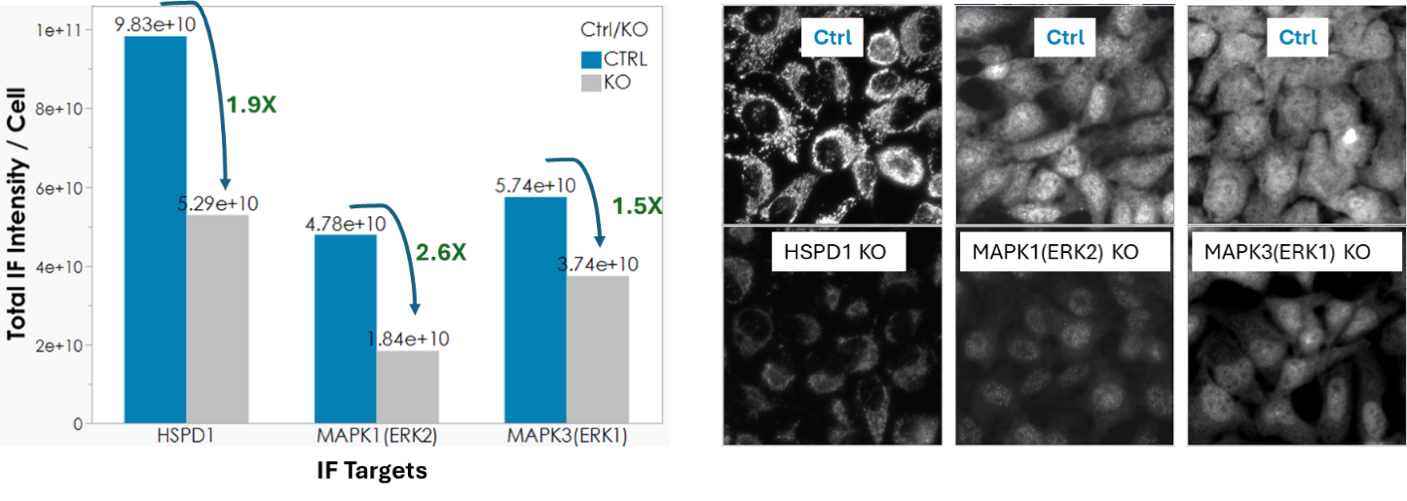

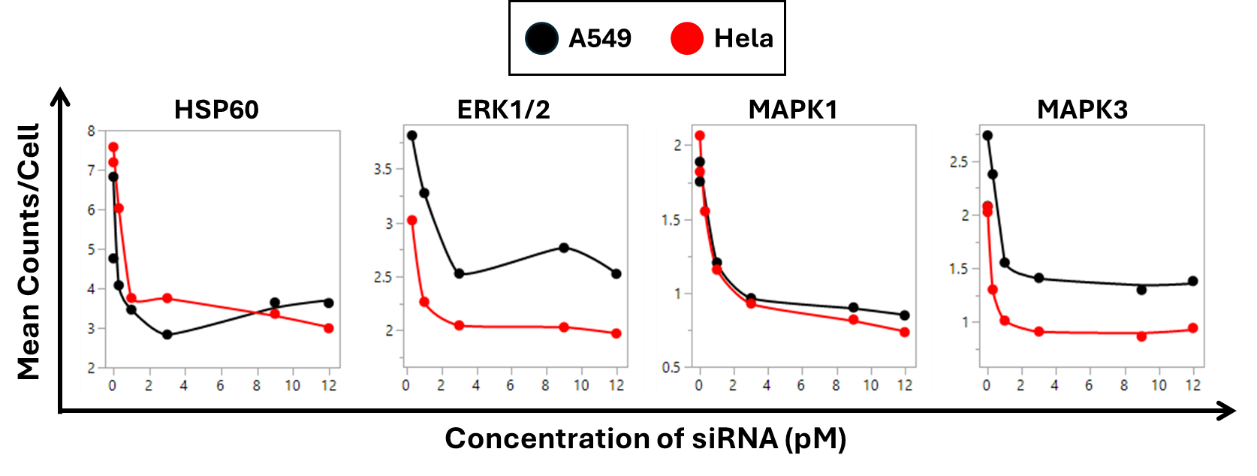

**Supplemental Figure 4: siRNA Knockdown Validation by Immunofluorescence and Dose-Response**

Top: Immunofluorescence signal intensities for HSPD1, MAPK1 (ERK2), and MAPK3 (ERK1) following siRNA-mediated knockdown in HeLa cells. Left: Quantification of total signal intensity per cell shows consistent reductions in KO compared to control conditions. Right: Representative fluorescence images for each knockdown confirm reduced signal.

Bottom: Dose-response profiles for the same targets across increasing siRNA concentrations in both HeLa (black) and A549 (red) cells. Each plot shows mean counts per cell, demonstrating robust, saturating knockdown effects with minimal off-target signal.

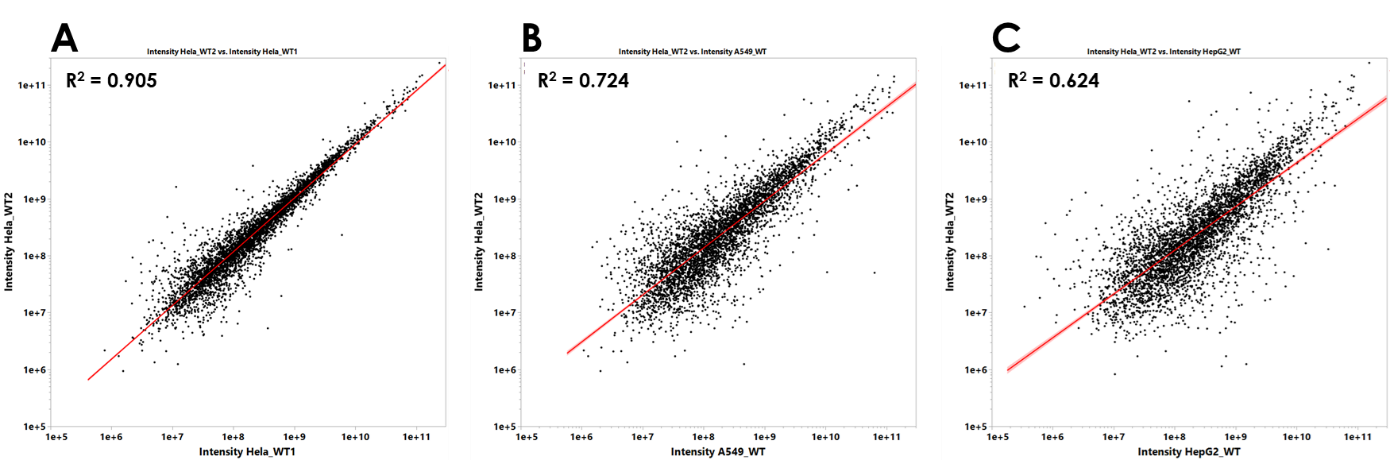

**Supplemental Figure 5: Label-Free Mass Spectrometry Comparison of Protein Expression Across Cell Types and Replicates**

(A) Technical replicates of HeLa cells (HeLa_WT1 vs HeLa_WT2) show a strong correlation in protein abundance (R² = 0.905), though considerable variability remains at lower abundance levels, limiting the confidence in differential expression calls except for the most significantly enriched proteins.

(B) Comparison of HeLa to A549 (R² = 0.724) and (C) HeLa to HepG2 (R² = 0.624) reveals greater divergence in protein profiles between cell types, as expected due to differential expression. However, the modest variability even between HeLa replicates highlights the limitations of label-free mass spectrometry in resolving subtle differences, suggesting that more quantitative methods such as isobaric labeling or targeted MS may be required to fully capture cell-type–specific proteomic changes.

| **Target** | **Colocalization (R) with commercial probe** |
| --- | --- |
| Cell Membrane | 0.71 ± 0.04 |
| Nucleus | 0.92 ± 0.01 |
| Mitochondria | 0.77 ± 0.05 |
| Actin | - |
| Golgi | 0.64 ± 0.06 |
| ER | 0.77 ± 0.03 |

*Error represents standard deviation across tiles

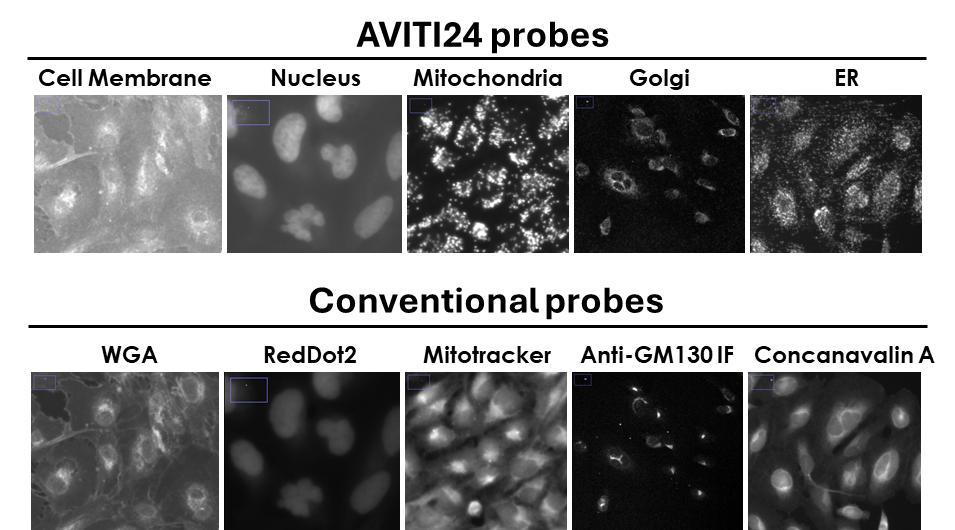

**Supplemental Figure 6: Colocalization between Teton Cell Paint and Commercial Markers.**

Representative images show the six Teton Cell Paint channels (top) alongside conventional fluorescent probes (bottom), including WGA (membrane), RedDot2 (nucleus), Mitotracker (mitochondria), anti-GM130 (Golgi), and anti-concanavalin (ER). Colocalization coefficients (R) between Teton probes and corresponding commercial markers are summarized in the table. Colocalization (R): Cell Membrane = 0.71 ± 0.04, Nucleus = 0.92 ± 0.01, Mitochondria = 0.77 ± 0.05, Golgi = 0.64 ± 0.06, ER = 0.77 ± 0.03.

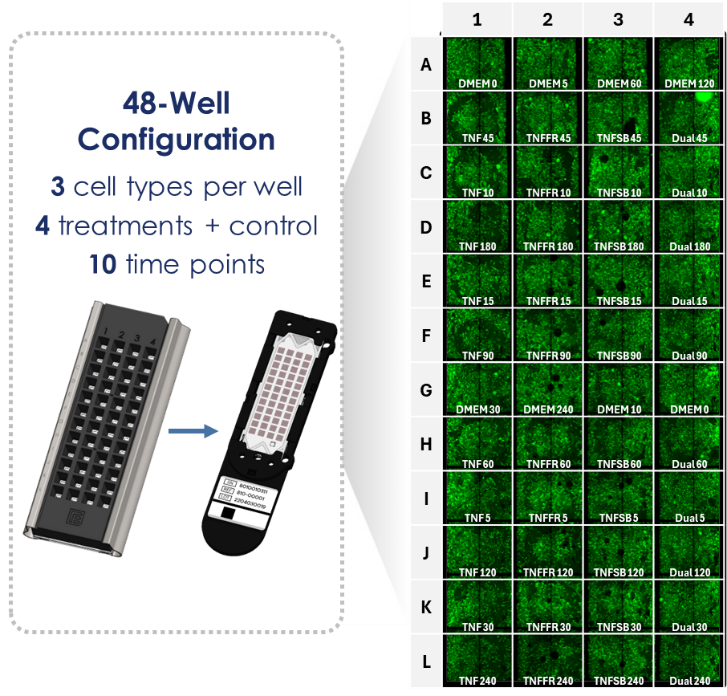

**Supplemental Figure 7: 48-well experimental layout for TNFα stimulation assay.**

Each well contains a co-culture of three cell types (HeLa, A549, and HepG2), with experimental conditions spanning four treatments (TNFα alone, TNFα + SB202190, TNFα + FR180204, TNFα + both inhibitors) and a control (DMEM). Timepoints range from 0 to 240 minutes across ten intervals. The right panel displays representative membrane staining (green) for each well, confirming even seeding across the flowcell.

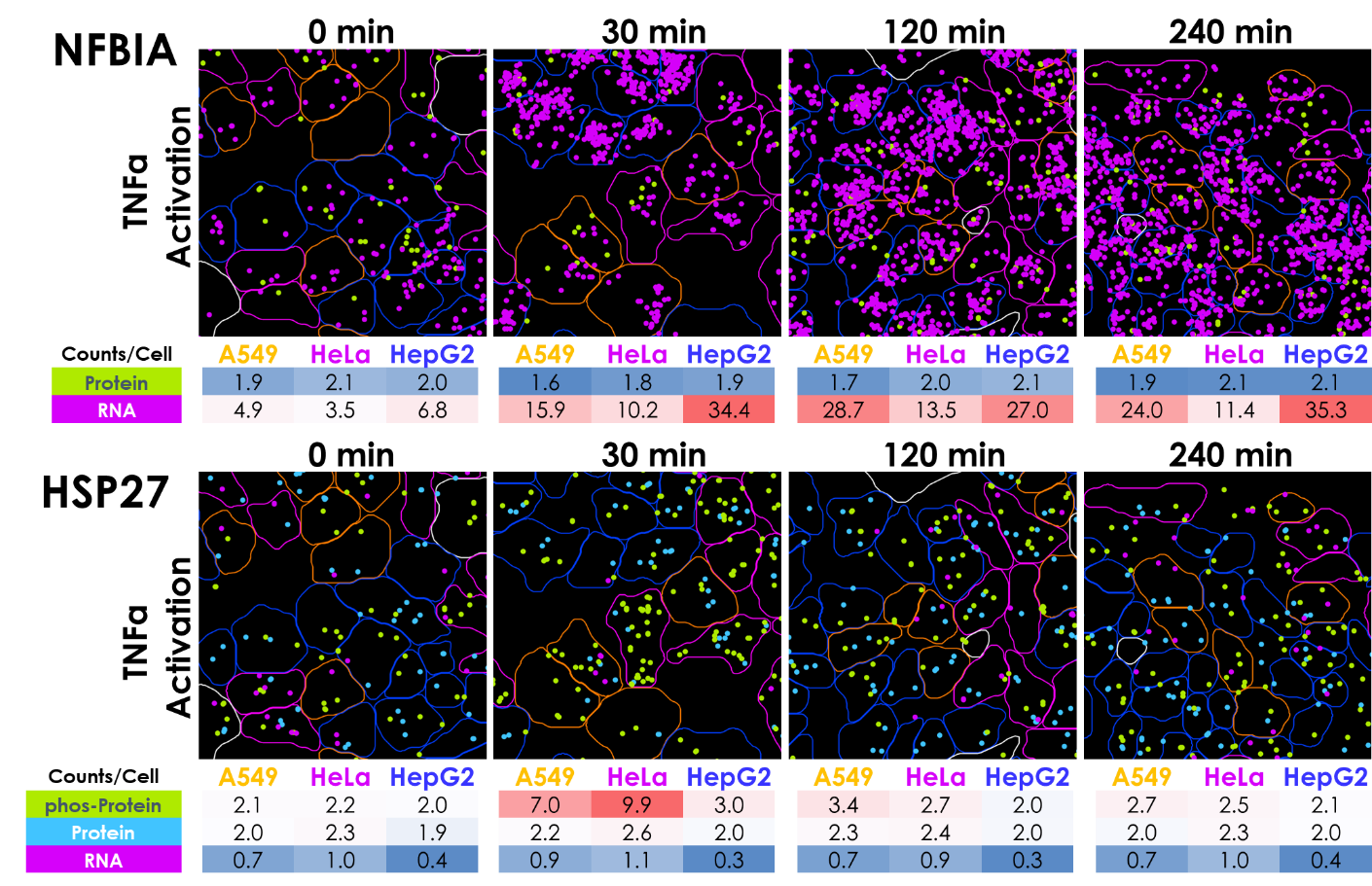

**Supplemental Figure 8: Single-Cell Multiomic Visualization of RNA, Protein, and Phosphoprotein Targets**

Representative images acquired on AVITI24 showing per-cell localization of RNA (magenta), protein (green), and phospho-protein (yellow) signals across four TNFα treatment timepoints. Each dot represents a polony assigned to a specific barcode. Cell boundaries are outlined by assigned cell type: A549 (orange), HeLa (magenta), HepG2 (blue), and filtered/unassigned (white). RNA and protein dynamics are shown for NFKBIA and HSP27, with mean counts per cell summarized in heatmaps below each panel. These images highlight distinct cell type–specific responses in target abundance and modality-specific regulation.

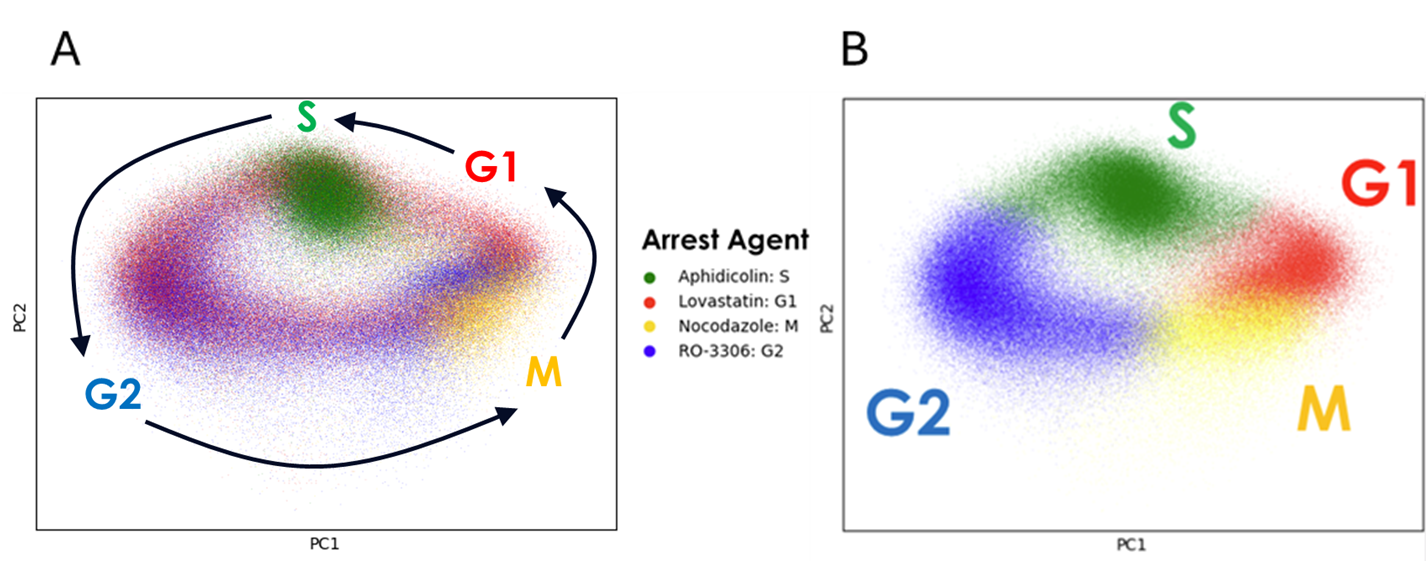

**Supplemental Figure 9: Cell cycle classifier training.**

(A) RNA features from two AVITI24 runs are projected onto the first two principal components, forming an annular structure characteristic of cell cycle progression. Cells were treated with different synchronization agents to arrest them in S, G1, G2, or M phases. Each dot represents a single cell, colored by the applied arrest agent. (B) Unsupervised clustering of all cells in (A) recapitulates cell cycle states, enabling assignment of PC clusters to canonical phases for downstream classification.

**Supplemental Figure 10: Temporal Dynamics of TNFα-Induced Multimodal Signaling Responses**

Time-lapse visualization of single-cell RNA and protein responses to TNFα stimulation and kinase inhibition across 10 time points (5–240 min) in HeLa, A549, and HepG2 cells. The GIF cycles through dynamic changes in expression levels for select RNA and protein targets, illustrating the temporal evolution of pathway activation and regulatory feedback. Each frame represents a different time point, showing consistent multimodal responses across treatment conditions and enabling resolution of transient signaling events at single-cell resolution. Targets include canonical stress and apoptotic regulators such as NFKBIA, BBC3, phospho-HSP27, and phospho-p38.

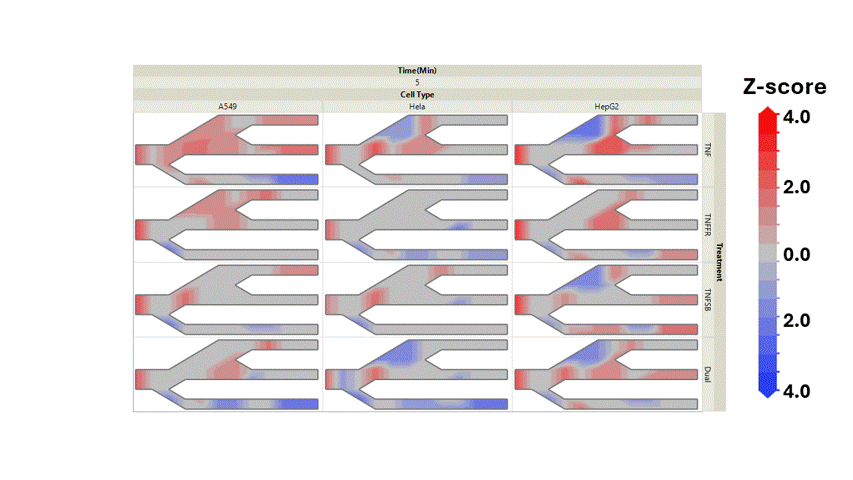

**DUSP**

**p38**

**ikBa**

**CyclinD1**

**HSP27**

**Bcl2A1**

**DUSP**

**p38**

**ikBa**

**CyclinD1**

**HSP27**

**Bcl2A1**

**DUSP**

**p38**

**ikBa**

**CyclinD1**

**HSP27**

**Bcl2A1**

**DUSP**

**p38**

**ikBa**

**CyclinD1**

**HSP27**

**Bcl2A1**

**DUSP**

**p38**

**ikBa**

**CyclinD1**

**HSP27**

**Bcl2A1**

**DUSP**

**p38**

**ikBa**

**CyclinD1**

**HSP27**

**Bcl2A1**

**DUSP**

**p38**

**ikBa**

**CyclinD1**

**HSP27**

**Bcl2A1**

**DUSP**

**p38**

**ikBa**

**CyclinD1**

**HSP27**

**Bcl2A1**

**DUSP**

**p38**

**ikBa**

**CyclinD1**

**HSP27**

**Bcl2A1**

**DUSP**

**p38**

**ikBa**

**CyclinD1**

**HSP27**

**Bcl2A1**

**DUSP**

**p38**

**ikBa**

**CyclinD1**

**HSP27**

**Bcl2A1**

**DUSP**

**p38**

**ikBa**

**CyclinD1**

**HSP27**

**Bcl2A1**
